# Treatment options for Chagas Disease: a systematic review and meta-analysis applied to the preclinical studies using animal models

**DOI:** 10.1101/2024.04.17.589953

**Authors:** Laura Yesenia Machaca-Luque, Mayron Antonio Candia-Puma, Brychs Milagros Roque-Pumahuanca, Haruna Luz Barazorda-Ccahuana, Luis Daniel Goyzueta-Mamani, Alexsandro Sobreira Galdino, Ricardo Andrez Machado-de-Ávila, Rodolfo Cordeiro Giunchetti, Eduardo Antonio Ferraz Coelho, Miguel Angel Chávez-Fumagalli

**Author notes:** These authors contributed equally to this work (Laura Yesenia Machaca-Luque, Mayron Antonio Candia-Puma).

## Abstract

Chagas disease (CD) is a neglected tropical disease endemic to Latin America and has emerged as a global health concern due to the migration of infected individuals. With its epidemiological complexity, difficulty in obtaining appropriate diagnoses, and poor treatment, the search for novel therapeutic options remains. In this context, we conducted a systematic review and meta-analysis of preclinical studies employing animal models to verify the progress in CD treatment. We searched the PubMed database for CD treatment studies published between 1990 and 2023, adhering to the PRISMA guidelines. Twelve papers met the inclusion criteria. The findings indicate that the fifteen treatment alternatives examined, mainly between 2010 and 2014, demonstrated efficacy in experimental CD models, evidenced by significant parasitemia reduction. Bis-triazole DO870 and VNI were effective in the acute and chronic phases, respectively. However, of these emerging therapies, only posaconazole and fexinidazole have progressed to clinical trials, yielding unsatisfactory outcomes as CD monotherapies. This meta-analysis highlights the existence of promising new drug candidates for CD treatment, but most remain in the preclinical stages. Those that reached clinical trials did not demonstrate optimal results, underscoring the ongoing challenges in CD therapy. Collaborative efforts among the academic community, pharmaceutical industries, funding agencies, and government agencies are urgently needed to accelerate the development of more effective medications against CD.

## Introduction

Chagas Disease, caused by the protozoan parasite *Trypanosoma cruzi*, is a neglected tropical disease endemic to Latin America, predominantly impacting rural and impoverished communities (1). Worldwide, the disease affects an estimated 6 to 7 million people, with 10,000 deaths each year (2). Additionally, there are emerging reports indicating the spread of Chagas Disease to non-endemic countries, fueled by factors such as human migration, travel patterns, globalization, and climate change, thereby introducing new complexities to disease management and surveillance efforts (3). The disease presents two distinct clinical phases: acute and chronic. Acute CD is often characterized by nonspecific symptoms such as fever, malaise, and local edema at the site of parasite entry (4), while chronic infection potentially leads to serious cardiac and gastrointestinal problems such as megaesophagus, megacolon, arrhythmias, and cardiomyopathy. All of these complications further increase the risk of sickness and death (5). The pathogenesis of CD is a complex process, with numerous factors affecting its transmission and spread (e.g., low disease occurrence in dogs or stray animals). To effectively combat this disease after so many years, many factors, including poor sanitation, shared housing, and what is known as "epidemiological blind spots," must be properly taken into consideration (6,7). Chronic infections can remain asymptomatic for years, only becoming aware whenever patients are infected even further. This makes infection control and patient care difficult problems indeed. Thus, there is an urgent need for new creative solutions to protect us from what constitutes an ongoing public health challenge (8).

The current therapeutic options for CD mostly consist of two nitroheterocyclic compounds, such as benznidazole and nifurtimox, developed in the late 1960s and early 1970s, respectively (9,10). Despite their decades-long employment, the medications offer limited efficacy, especially in the chronic phase of the disease, and are associated with significant side effects that often necessitate extensive treatment courses (11). The management of CD is further plagued by difficulties stemming from low patient adherence, drug outdating, drug resistance, and therapeutic failure associated with existing therapies (8,11). The development of new drugs for CD has been hampered by various factors, including the lack of financial incentives for pharmaceutical companies to develop drugs for a disease that mainly affects impoverished populations in low-resource settings and the complex biology of the parasite and host immune responses (12,13). Not surprisingly, there is an unmet need for innovative research initiatives and more effective collaborations that will address the numerous therapeutic challenges posed by CD and improve the number of patients with this disease (14,15).

Recently, fundamental research, particularly in the preclinical stage, has been the focal point of efforts aimed at developing new treatment strategies for CD. These endeavors predominantly utilize techniques derived from *in vitro* or animal investigations (10,16). Notably, animal model-based preclinical research has emerged as indispensable for evaluating novel CD treatment approaches (17). Such studies facilitate the exploration of fundamental mechanisms of action and pharmacokinetic characteristics, while also providing valuable insights into the safety and efficacy of innovative therapies (18). In light of these advancements, the current study aims to conduct a systematic review and meta-analysis of preclinical research utilizing animal models, to assess the efficacy of treatment options for CD.

## Methods

### Study protocol

This systematic review was conducted following the Preferred Reporting Items for Systematic Reviews and Meta-Analyses (PRISMA) statement (Table S1) (19). With registration number INPLASY202430101, the protocol for this systematic review was registered on the International Platform of Registered Systematic Review and Meta-analysis Protocols (INPLASY) website. The entire protocol is accessible at inplasy.com (https://inplasy.com/inplasy-2024-3-0101/).

### Information sources and search strategy

The literature was searched for phrases about CD therapy using the MeSH (Medical Subject Headings) term *"Chagas Disease".* Using the VOSviewer program (version 1.6.20), the findings were shown in a network diagram showing the co-occurrence of MeSH keywords (20). We looked at clusters in the network map to choose phrases associated with CD therapy. Furthermore, a second round of searches was carried out by linking each MeSH term identified in the cluster analysis with the MeSH terms *"Chagas Disease"* and *"Treatment Outcome",* which relate to assessing the outcomes of interventions used to combat diseases and determining their efficacy (21), For the years 1990–2023, records were obtained from the bibliographic database PubMed (https://pubmed.ncbi.nlm.nih.gov/, last accessed 24 May 2023).

### Selection criteria and data extraction

The procedure for selecting studies for this review comprised three separate phases. During the initial identification phase, only animal studies published between 1990 and 2023 were taken into account. Duplicate articles, non-English publications, reviews, and meta-analyses were excluded at this stage. The subsequent screening phase involved checking the titles and abstracts of the identified articles, and in the eligibility/qualification phase, full-text studies highly relevant to the research question were retrieved, specifically focusing on treatment options for CD. Data on the type of compound used for treatment, dosage, duration of treatment, total sample size, number and species of experimental animals infected, phase of CD, *T. cruzi* strain, sample type, and description of the controls were extracted from each of the chosen studies. Studies with insufficient data were excluded. In contrast, those who assessed the effectiveness of therapy by disclosing parasitemia data were retained, considering a 70% decrease in parasitemia as a positive treatment for both the treated and control groups. Options for treatment that included a drug in addition to a conventional CD medication, like nifurtimox or benznidazole, weren’t included. Drugs used in monotherapy were emphasized to confirm which ones show higher effectiveness. WebPlotDigitizer was used for data analysis when the information was shown graphically (22,23). WebPlotDigitizer is a semi-automatic tool that lets you manually plot two-dimensional charts and extract numerical data. It is free online (https://automeris.io/WebPlotDigitizer/) or as desktop software that may be downloaded (22). This instrument has been employed in several systematic reviews and meta-analyses, encompassing therapy efficacy assessment (24–26). L.Y.M.-L. carried out the data extraction, and M.A.C.-P. Independently checked it. If there were any differences, M.A.C.-F. was consulted and discussed.

### Statistical analysis

Results were entered into a Microsoft Excel (version 2108, Microsoft Corporation, Redmond, WA, USA) spreadsheet and analyzed in the R programming environment (version 4.2.3) using the "*metafor*" package https://www.metafor-project.org/doku.php/metafor (accessed on 21 February 2024) (27). Plotting the synthesis findings and estimating a random-effects model are just a few of the numerous tasks that may be performed by the user with the help of the “*metafor*” package (27,28). The number of treated participants who tested positive or negative for CD (t_pos_ and t_neg_, respectively), and the corresponding number of untreated patients (control) who tested positive or negative for CD (c_pos_ and c_neg_, respectively), were evaluated independently for each therapeutic option available with the *“metafor”* package”.

A Random Effects Model (RE model) was utilized in the meta-analysis process. According to this model, every study has a unique true effect that varies depending on the differences in participants, interventions, or circumstances between studies (29). The Q value (Q), the I-squared (I^2^), the tau-squared (T^2^), and the risk ratio (RR) were also computed. One indicator of heterogeneity among the studies in the meta-analysis is the Q. The sum of the squares representing the discrepancies between the weighted overall effect and the observed effects are used to compute it. A high Q score (30) indicates higher levels of study heterogeneity.

The overall percentage of variance in the estimated effects that results from heterogeneity between studies as opposed to random variation is measured by the I^2^ statistic. It is represented as a percentage and computed as (Q - df) / Q (31). Excessive I^2^ values (above 50%) suggest significant variability among the studies (32). In a random effects model, the variation between studies is represented by the T^2^. Beyond sample error, it evaluates the real dispersion of effects (33). More significant variability between trials that cannot be attributed to chance is indicated by a higher T^2^ score (34). As the RR of the event happening in the exposed or treated group compared to the unexposed or untreated group, the RR measures the connection between an exposure or treatment and an outcome of interest (35). Compared to the reference group, an exposure or treatment is linked to a higher outcome risk if the relative risk (RR) is more significant than 1. Conversely, a lower risk is suggested when the RR is less than 1. If the relative risk (RR) is 1, then there is no variation in risk among the groups. Confidence intervals are a useful tool for assessing the accuracy of the estimate, in addition to the RR value (36). For all computations, a 99% confidence level was used, with a 0.1 continuity adjustment applied as necessary.

## Results

### Data sources and study selection

In this work, we performed a systematic review and meta-analysis to evaluate the effectiveness of many treatment options for lowering parasitemia. We considered both established therapies and recently developed, potentially effective medicines for CD. **Figure 1** shows a flowchart of the study approach. A search for the MeSH term *“Chagas Disease”* AND *“Treatment Outcome”* was performed in the PubMed database and a MeSH term co-occurrence network map was developed. Through the search, 323 scientific publications were published between 1990 and 2023. A network map with 1,020 keywords was produced, with the minimal number of keyword occurrences set at five (**Figure 2**). Three major clusters were found to have formed during the network map study. Terms including *"nitroimidazoles”*, *"nifurtimox"*, *"benznidazole"*, and *"antiprotozoal agents"* were found in the cluster about the therapy of CD (highlighted in green). There were other similar denominators, including phrases like *"treatment outcome"*, *"humans"*, *"Chagas disease"*, *"female"*, *"male"*, and "*Trypanosoma cruzi*" (**Figure 2**).

**Figure 1.**
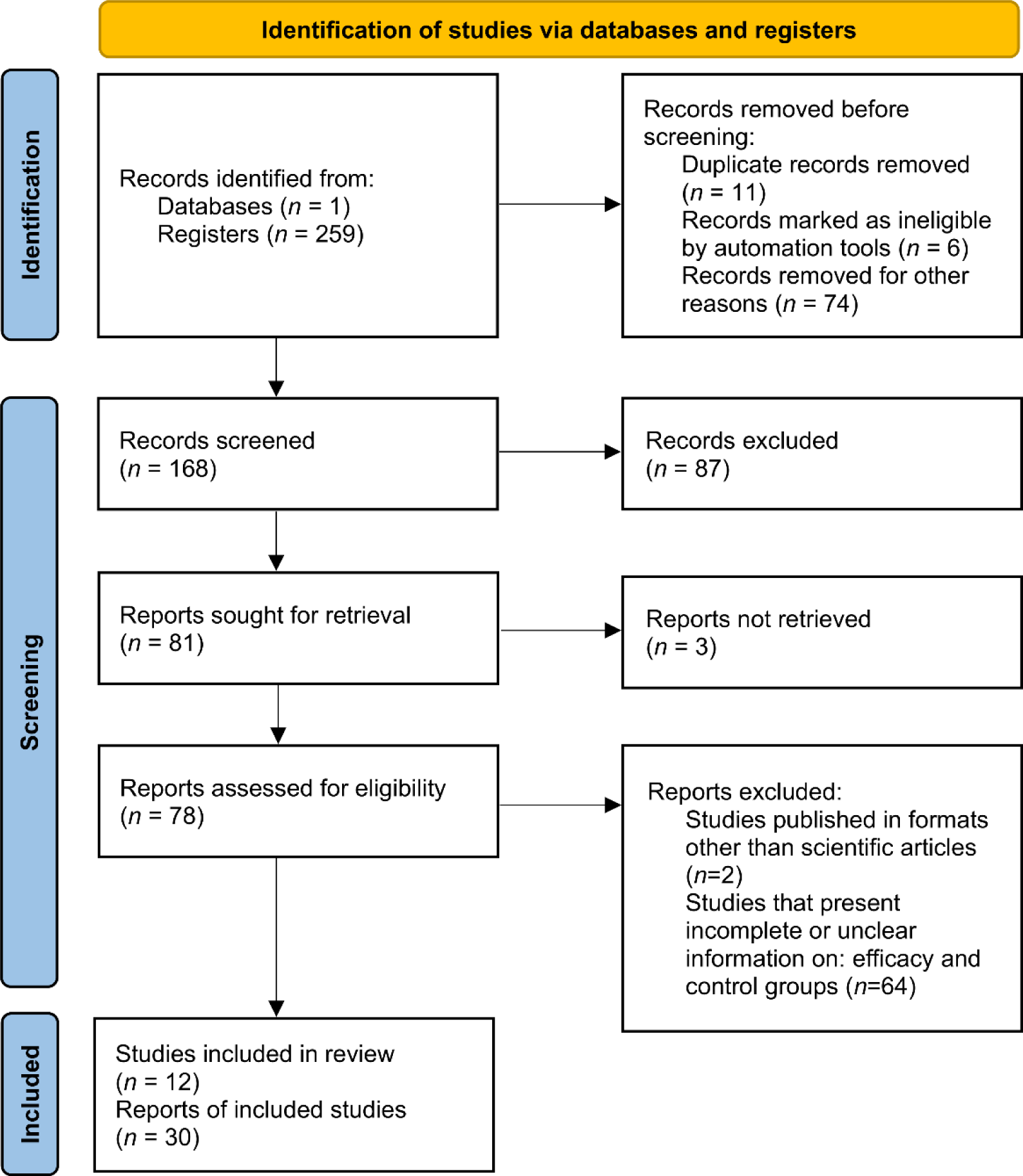
A flowchart representing the study selection process’s meta-analysis and systematic review.

**Figure 2.**
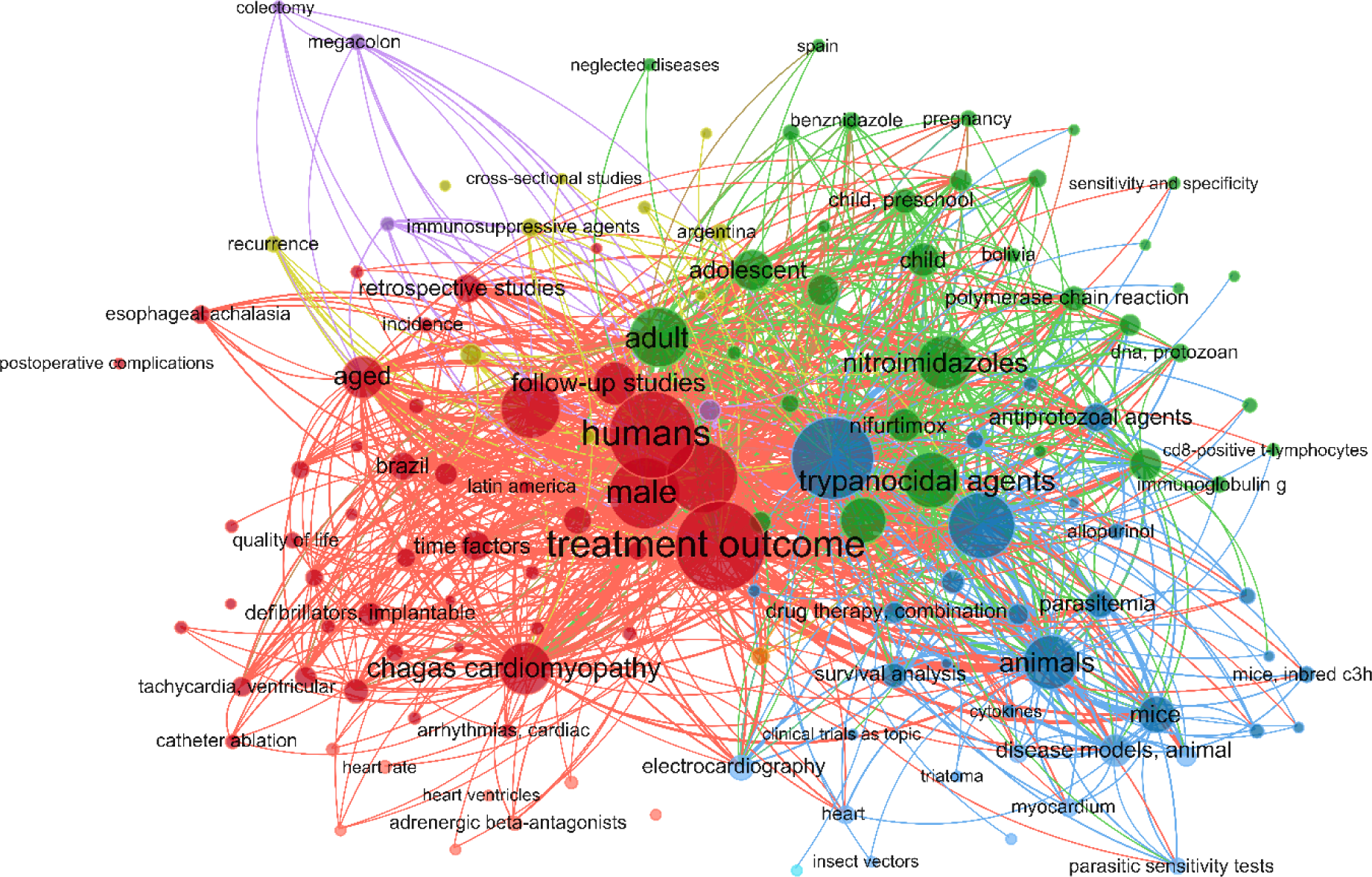
A network diagram was made with VOSviewer using the PubMed database to represent the treatment result of CD, based on the incidence of MeSH terms in a selected set of publications.

The terms found in the first analysis were used to perform a second search in the PubMed database. After the new phrases were associated with *"Chagas Disease"* and *"Treatment Outcome",* the following new search strings were generated: (Chagas Disease[MeSH Terms]) AND (Treatment Outcome[MeSH Terms]) AND (nifurtimox[MeSH Terms]), (Chagas Disease[MeSH Terms]) AND (Treatment Outcome[MeSH Terms]) AND (nitroimidazoles[MeSH Terms]), and (Chagas Disease[MeSH Terms]) AND (Treatment Outcome[MeSH Terms]) AND (Therapeutics[MeSH Terms]) for commonly used treatments (nifurtimox and benznidazole) as well as recently developed treatments against CD.

There were 40, 107, and 112 studies chosen for the first, second, and third search strings, respectively. We excluded 91, 90, and 66 articles in the identification, screening, and eligibility phases, respectively, based on the three-step selection criteria that we employed. This resulted in the selection of 12 studies for the meta-analysis, one published in 2001, and the others between 2010 and 2014, several of which discussed various treatment choices; as a result, 30 publications in total were included in the investigation. In these investigations, the following novel chemical compounds were examined: Posaconazole, AmBisome^®^, Cyclopalladated complex 7a, Fexinidazole, Psilostachyin A, Cynaropicrin, Reversible cruzipain inhibitors Cz007 and Cz008, dehydroepiandrosterone-sulfate, VNI, (−)−hinokinin-loaded microparticles, allopurinol, clomipramine, GW788388, and Bis-triazole D0870 (**Table 1**). The most significant number of studies utilizing animal models to investigate potential novel treatments for CD have been conducted in Brazil. In addition, albeit to a lesser degree, studies on this subject have also been done in several other nations, including Argentina, Canada, France, and the United States (**Figure 3**).

**Figure 3.**
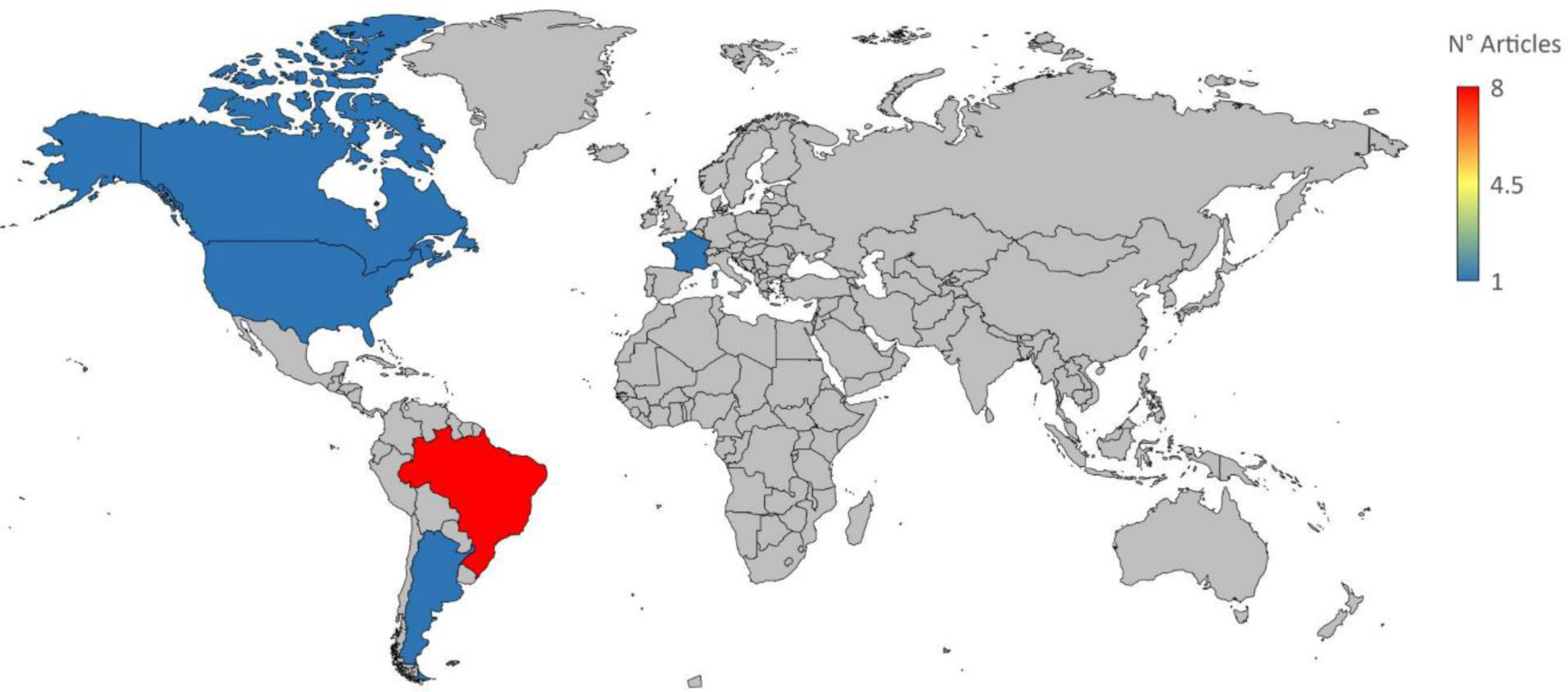
A demographic breakdown of the international research on novel CD therapy options that were included in the meta-analysis (numbers ranging from lower blue to upper red).

**Table 1.**
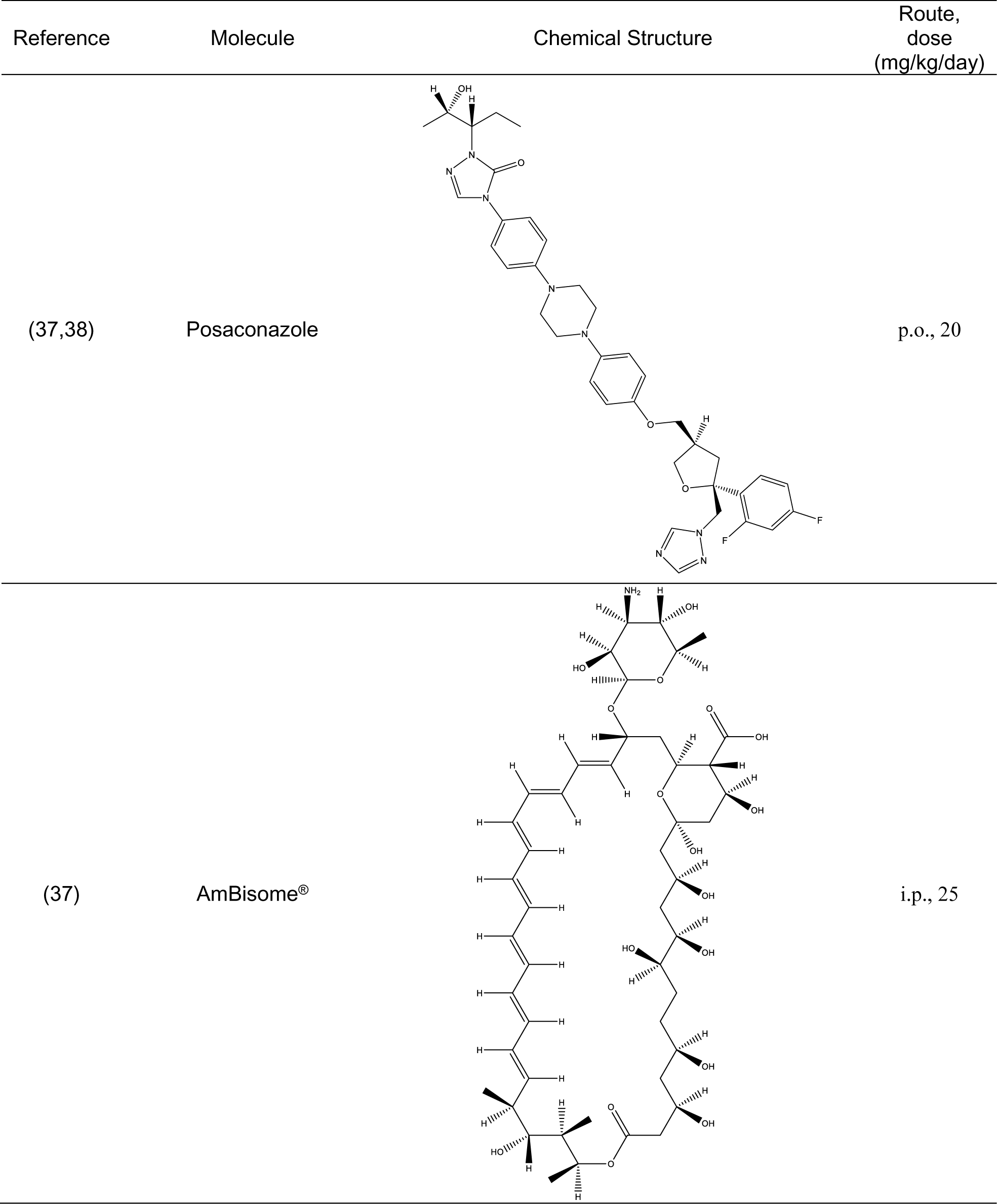
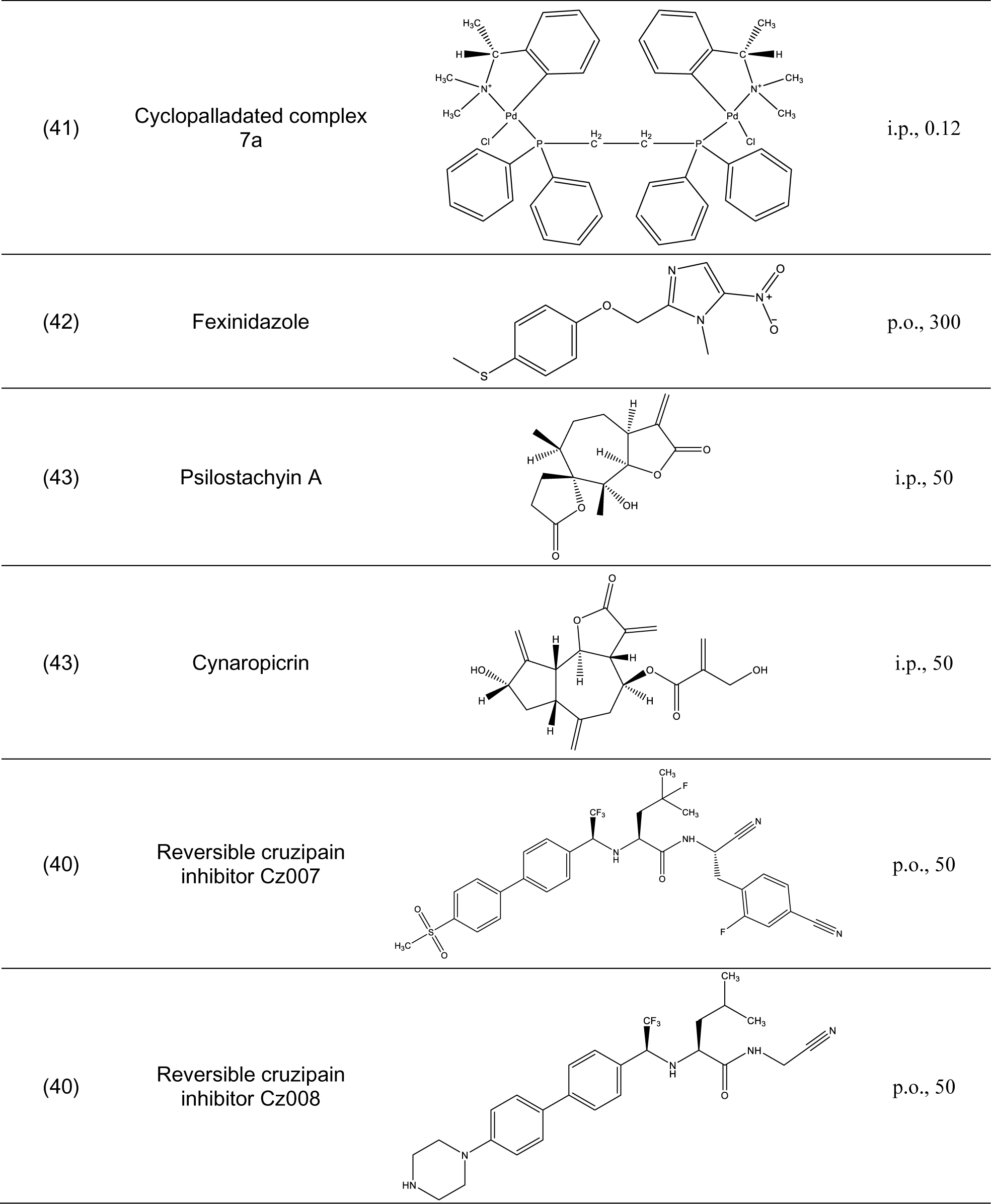
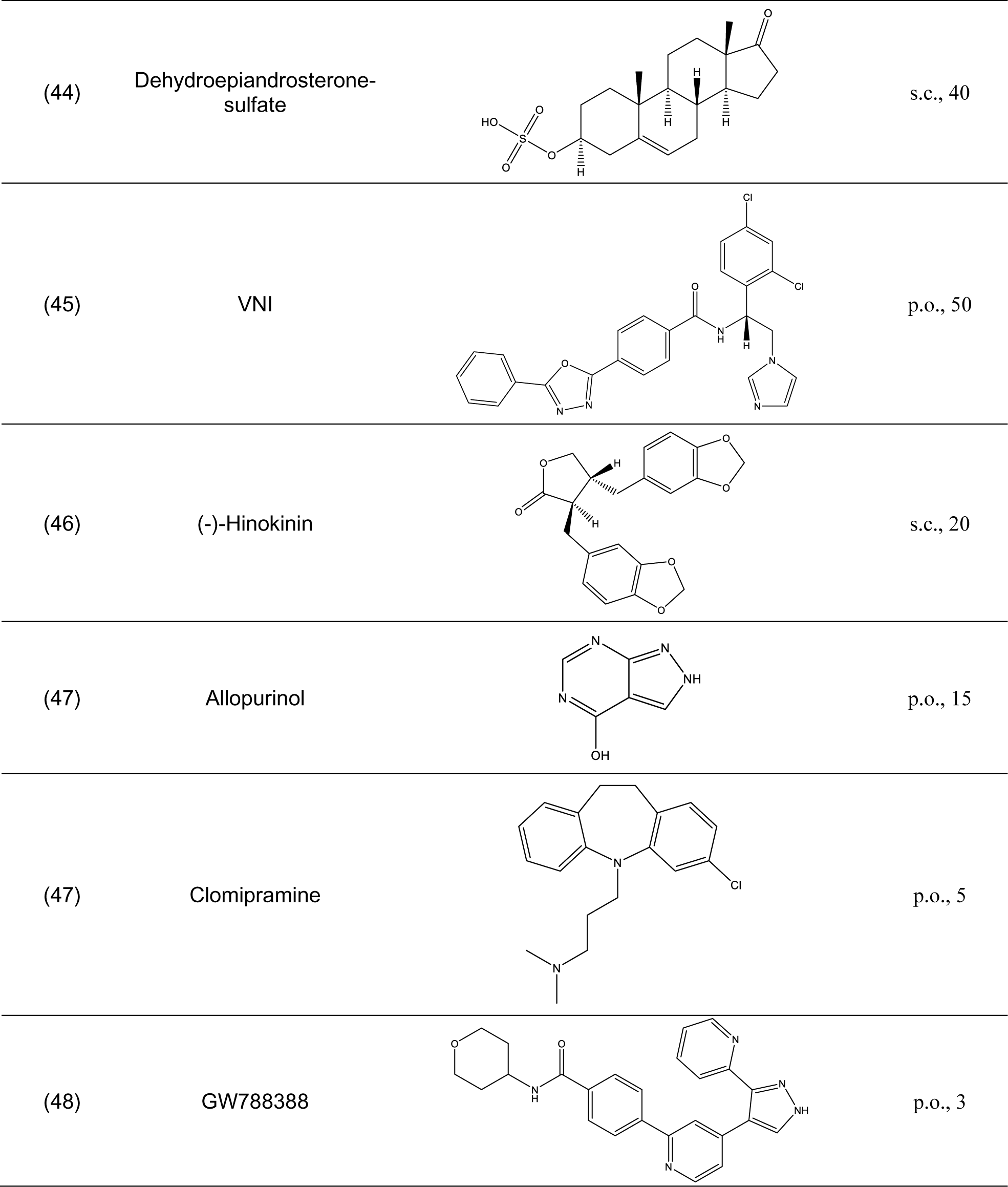
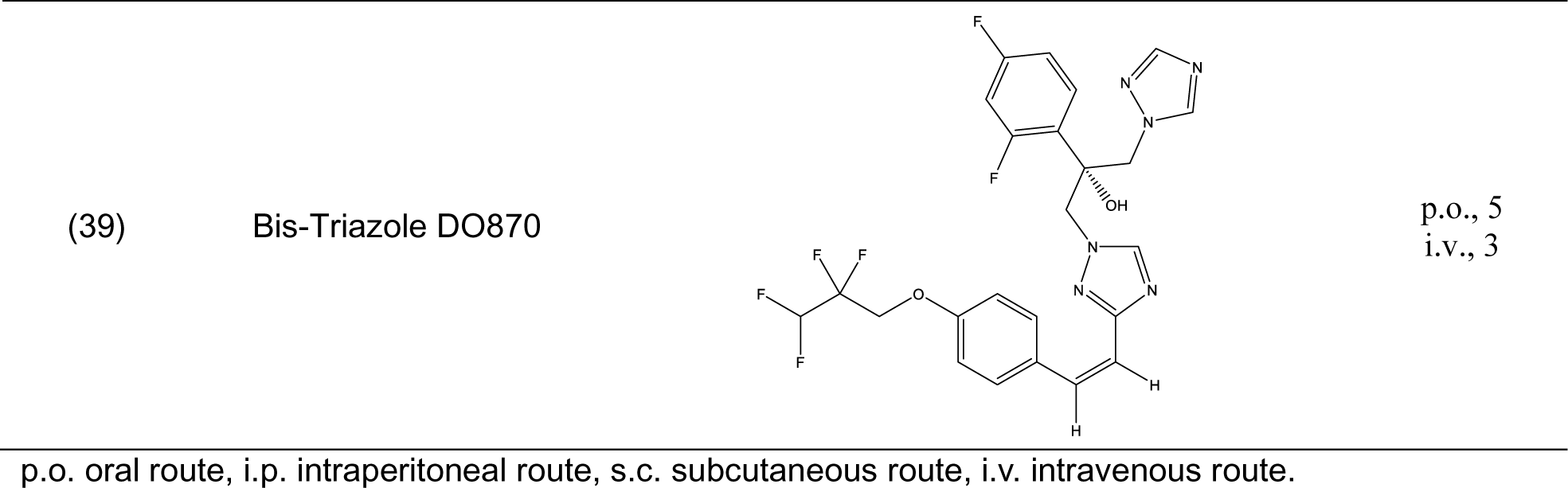
Chemical compounds evaluated in preclinical studies using animal models of CD.

### Meta-analysis of the treatment options for CD CD phase

The acute phase of the disease and its treatment options posaconazole, AmBisome®, Cyclopalladated complex 7a, Fexinidazole, Psilostachyin A, Cynaropicrin, Reversible cruzipain inhibitors Cz007 and Cz008, dehydroepiandrosterone-sulfate, VNI, (−)−hinokinin-loaded microparticles, allopurinol, clomipramine, GW788388, and Bis-triazole D0870 were also covered in twelve studies (37–48) that included a total of 741 animals. As **Figure 4** shows, the treatment alternatives, particularly Bis-triazole D0870, proved to be effective compared to the control groups. (RR = 8.22, 95% CI [4.80, 14.06]). The tests showed that there was minimal heterogeneity (Q (df = 24) = 14.31, p = 0.94; I^2^ = 0.0%; T^2^ = 0.0%)). Two studies (37,45) with a total of 138 animals were examined that dealt with the chronic phase of the disease and its treatment choices (VNI, Benznidazole, Nifurtimox, Posaconazole AmBisome®). The treatment alternatives were effective compared to the control groups, especially VNI, as **Figure 4** illustrates (RR = 11.29, 95% CI [3.32, 38.36]). There was minimal heterogeneity, according to the tests (Q (df = 4) = 0.33, p = 0.99; I^2^ = 0.0%; T^2^ = 0.0%)).

**Figure 4.**
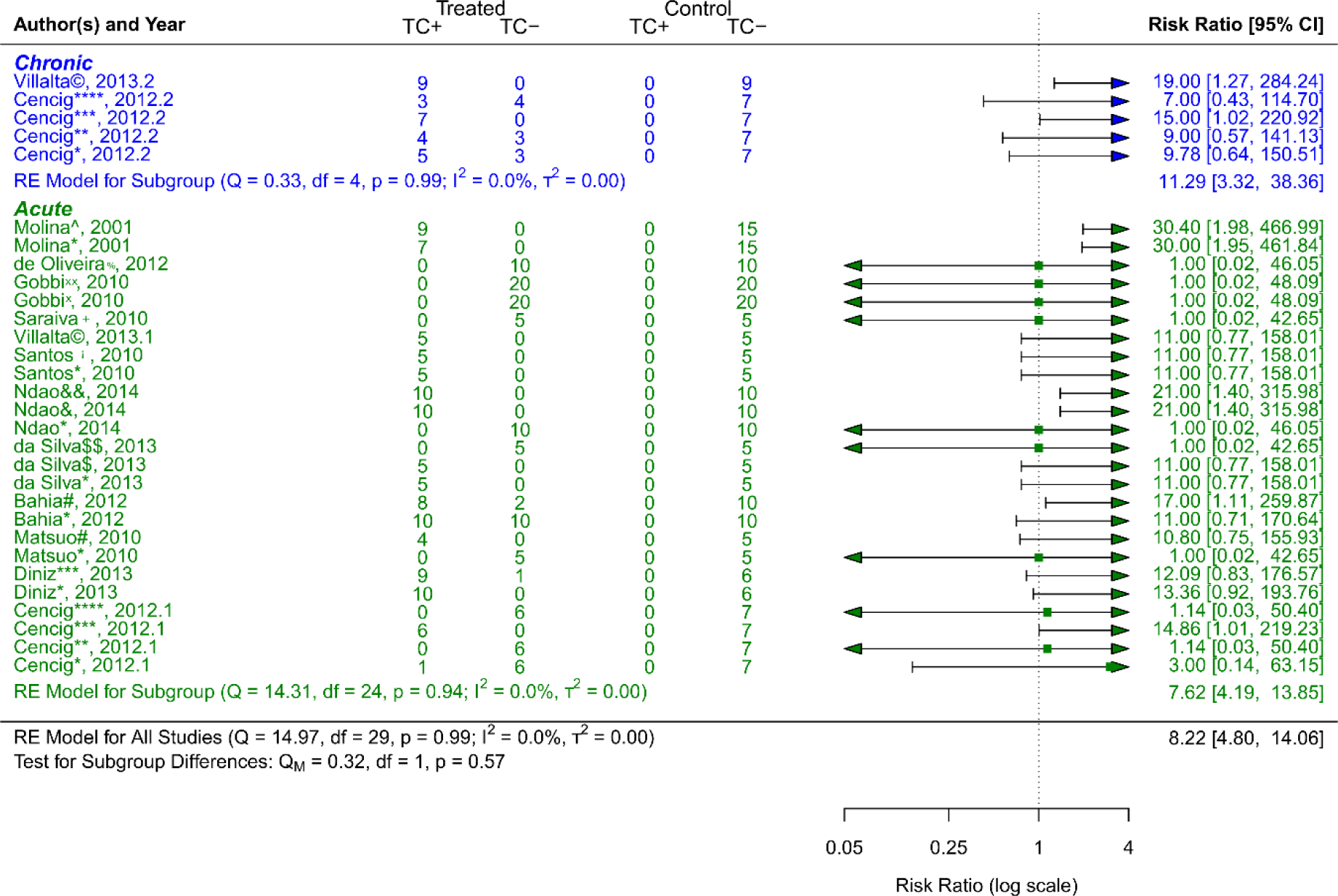
Forest plot showing a comparison of efficacy (estimated average response rates RR) between treated groups and control groups according to CD phase (acute and chronic). Symbol meaning: * Benznidazole, ** Nifurtimox, *** Posaconazole, **** AmBisome^®^, # Cyclopalladated complex 7ª, ## Fexinidazole, $ Psilostachyin A, $$ Cynaropicrin, & Reversible cruzain inhibitors Cz007, && Reversible cruzain inhibitors Cz008, ¡ Dehydroepiandrosterone sulfate, © VNI, + (−)−hinokinin-loaded microparticles, x Allopurinol, xx Clomipramine, % GW788388, and ^ Bis-triazole D0870. Error bars represent 95% CI. The square shapes represent the estimated RR. The vertical dashed line represents the no-effect line.

### Experimental animal models - strains

The scientific articles were categorized into smaller groups based on the type of animal model that received a *T. cruzi* inoculation. Only one study (44) employed Wistar strain rats. Other mice strains were employed in the other studies: Swiss (37,38,40,42,43), BALB/c (43,46,47), CD-1 (39,48), and BALB/cJ (37,41,45). As can be observed in Figure 5, CD−1 was the strain that responded to the treatment the best out of all the investigated strains. Its RR = 11.41, 95% CI [2.05, 63.36], indicates this. The tests showed that there was very little heterogeneity (Q (df = 2) = 1.94, p = 0.38; I^2^ = 0.0%; T^2^ = 0.0%)). On the other hand, the strain with the highest number of trials was the Swiss strain, which as shown in **Figure 5** with an RR = 8.02, 95% CI [3.49, 18.46]) also showed a substantial response to the various treatment methods (Q (df = 12) = 8.21, p = 0.77; I^2^ = 0.0%; T^2^ = 0.0%) showed low heterogenicity. With an RR = 6.58, 95% CI [2.32, 18.69], and negligible heterogenicity (Q (df = 7) = 2.74, p = 0.91; I^2^ = 0.0%; T^2^ = 0.0%), BALB/cJ was the strain that responded to the treatment alternatives the least (**Figure 5**).

**Figure 5.**
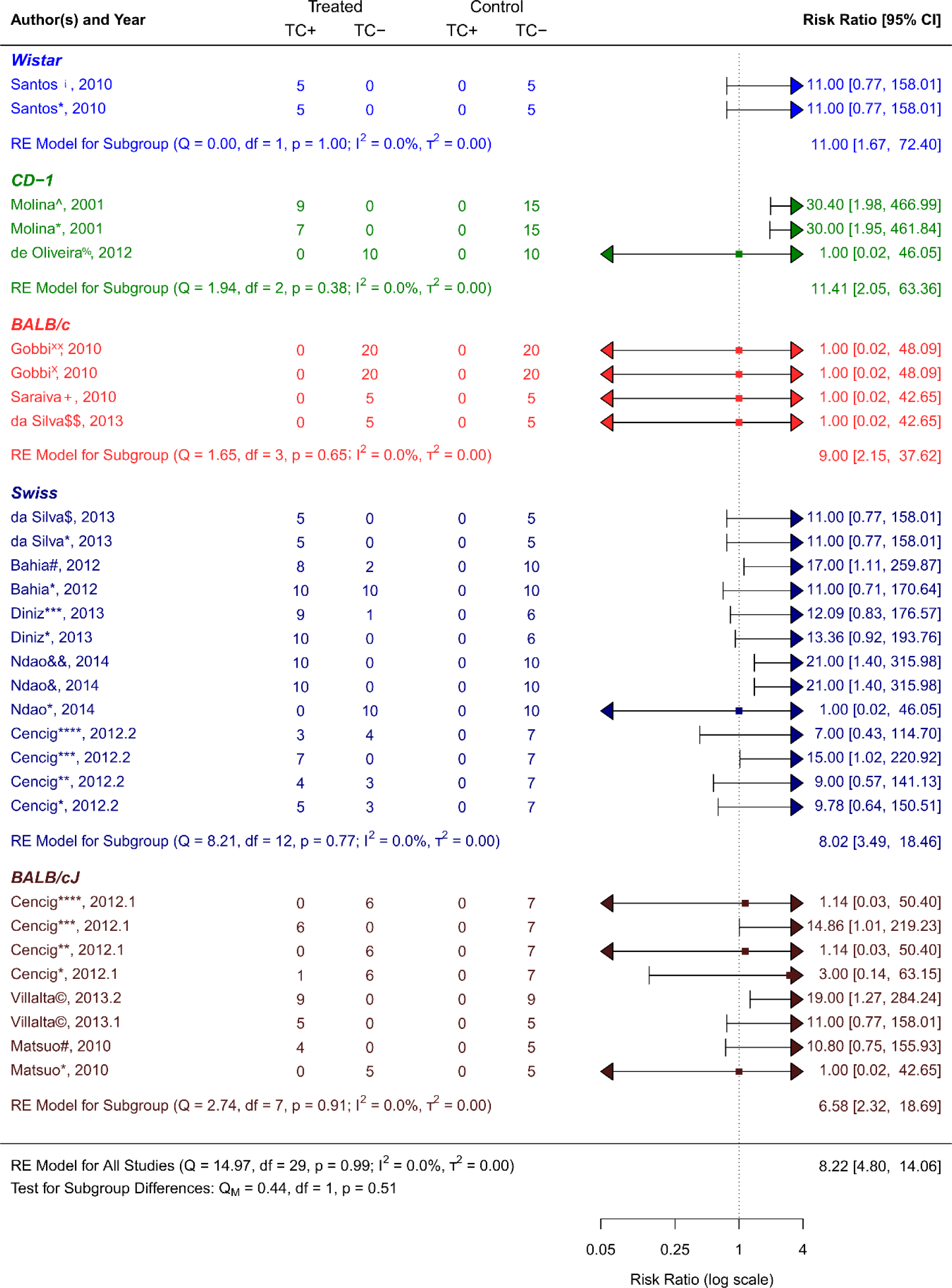
Forest plot showing a comparison of efficacy (estimated average response rates RR) between treated groups and control groups according to the strain of the animal (model Wistar, CD−1, BALB/c, Swiss, and BALB/cJ) used in the studies. Symbol meaning: * Benznidazole, ** Nifurtimox, *** Posaconazole, **** AmBisome®, # Cyclopalladated complex 7ª, ## Fexinidazole, $ Psilostachyin A, $$ Cynaropicrin, & Reversible cruzain inhibitors Cz007, && Reversible cruzain inhibitors Cz008, ¡ Dehydroepiandrosterone sulfate, © VNI, + (−)−hinokinin-loaded microparticles, x Allopurinol, xx Clomipramine, % GW788388, and ^ Bis-triazole D0870. Error bars represent 95% CI. The square shapes represent the estimated RR. The vertical dashed line represents the no-effect line.

### Experimental animal models - sex

The sex of the animal model was used to categorize the scientific studies. RR = 9.36, 95% CI [4.83, 18.12] was the result of 7 studies including female experimental animals (37–39,41,42,45,46); Q (df = 18) = 7.91, p = 0.98; I^2^ = 0.0%; T^2^ = 0.0%, indicated a low level of heterogenicity (**Figure 6**). The studies with the male sex of the experimental animals were 5 (40,43,44,47,48), showed a RR = 6.38, 95% CI [2.54, 16.04], with heterogenicity data of Q (df = 10) = 6.62, p = 0.76; I^2^ = 0.0%; T^2^ = 0.0% (**Figure 6**). While analyzing the results, it is clear that female experimental animals are more susceptible than male counterparts to the various CD therapy approaches.

**Figure 6.**
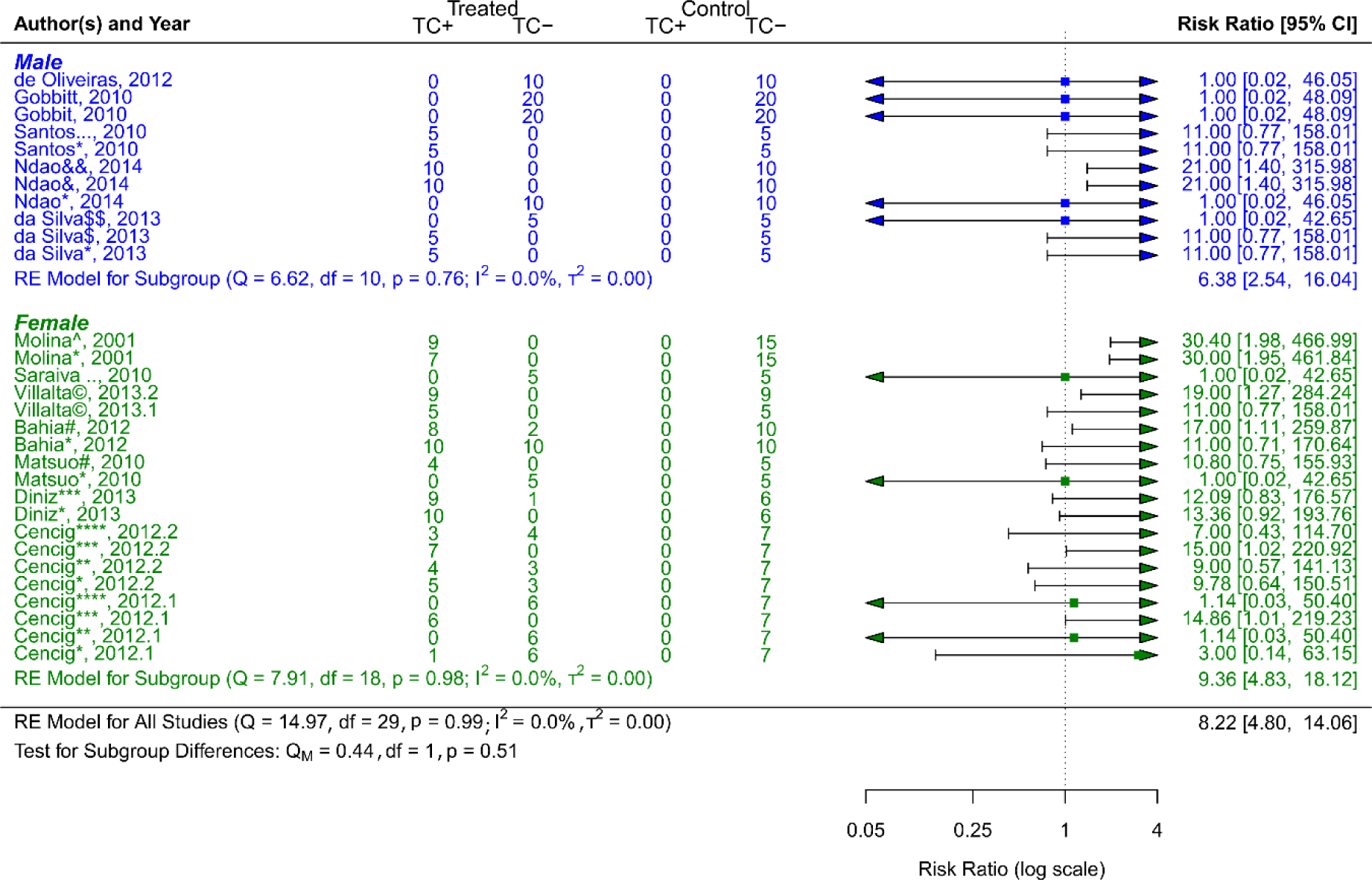
Forest plot showing a comparison of efficacy (estimated average response rates RR) between treated groups and control groups according to the sex of the animal model used in the studies. Symbol meaning: * Benznidazole, ** Nifurtimox, *** Posaconazole, **** AmBisome®, # Cyclopalladated complex 7ª, ## Fexinidazole, $ Psilostachyin A, $$ Cynaropicrin, & Reversible cruzain inhibitors Cz007, && Reversible cruzain inhibitors Cz008, ¡ Dehydroepiandrosterone sulfate, © VNI, + (−)−hinokinin-loaded microparticles, x Allopurinol, xx Clomipramine, % GW788388, and ^ Bis-triazole D0870. Error bars represent 95% CI. The square shapes represent the estimated RR. The vertical dashed line represents the no-effect line.

### Other research on CD treatments

In the final analysis of the scientific publications (49), one article about immunotherapy using DNA vaccines in mice was discovered. Immunotherapy boosts the body’s immune response, whereas conventional therapies utilize medications to treat a disease’s symptoms or underlying causes (50). This study was left out of the meta-analysis because of their disparate biological objectives, which make them hard to evaluate in terms of efficacy and safety.

## Discussion

### Summary of main findings

Most of the research that made up the meta-analysis was developed between 2010 and 2014. A complex and multidimensional junction of circumstances has impeded development in CD research, leading to an extended time of investigations on new therapeutic drugs. From a scientific standpoint, because of the disease-causing *T. cruzi*’s inherent genetic variety and capacity to elude the host immune system, its biological complexity has made it extremely difficult to pinpoint effective treatment targets (51–53). Moreover, the formidable challenge facing researchers in conducting extensive clinical investigations and acquiring the necessary resources to drive therapeutic innovation has been exacerbated by inadequate funding for CD-specific research and therapy development (54–56). Heightening this issue is the pharmaceutical industry’s limited interest in the subject, exacerbated by the meager financial prospects of medicines targeting illnesses primarily affecting low-income regions (57,58). The emergence of drug-resistant strains underscores the urgent need for alternative therapeutics, while regulatory hurdles associated with approving novel treatments for neglected diseases further compound the delays in progress (53,59–61). Collectively, these factors converge to create a daunting and intricate landscape that significantly hampers the development of novel CD treatments.

A persistent lack of funding dedicated to CD research and treatment development is a significant obstacle to therapeutic progress (54–56). The lack of financial resources limits researchers’ capacity to conduct robust clinical trials and acquire essential resources. Compounding this issue, the pharmaceutical industry shows limited interest due to the low commercial potential of medications targeting diseases prevalent in low-income regions (57,58). Furthermore, the emergence of drug-resistant strains emphasizes the urgency of finding alternative treatments, yet regulatory hurdles associated with approving novel therapeutics for neglected diseases further contribute to delays (53,59–61). Collectively, these factors create a complex and discouraging landscape that severely hinders the development of new CD treatments.

In Brazil, researchers conducted three-quarters of the studies that comprise this meta-analysis. This country is a natural hub for study in this field since it is one of the nations most impacted by CD, has a robust research infrastructure, and an epidemiological database (62,63). Research in the nation has also been enhanced by the presence of academic institutions, centers of excellence in tropical health, and financial and research resources (64). On the other hand, even though other Latin American nations are also impacted by CD, scientific research and the health system face structural and financial obstacles that may hinder study (65). Regarding North American countries like the United States of America and Canada, involvement in CD research may depend on the financing available for global health research initiatives and the interests of certain academics or groups (66,67). The lack of urgency in researching and developing therapies may be attributed to the disease’s low incidence in this area (67,68). Only one study from France was considered in the meta-analysis. However, it should be considered that, although non-endemic countries do not directly suffer from CD in their populations, they should consider globalization and human mobility, since these factors increase the possibility of transfer. of diseases to non-endemic countries (69–71). Beyond the moral need to alleviate patients’ suffering, the worldwide scope of CD necessitates a cooperative, international effort to solve the problems this neglected tropical disease presents.

The results of this study, bis-triazole DO870 and VNI have demonstrated promise as treatments for CD in the acute and chronic stages, respectively. The assessment of these substances’ safety and effectiveness in actual clinical settings is unknown, as no published clinical trials have been performed too far. Similarly, only two of the treatment options included in this meta-analysis—posaconazole and fexinidazole—have undergone clinical trial evaluation. Posaconazole’s clinical research shows that, although exhibiting trypanostatic action during therapy, it was ineffective in treating asymptomatic *T. cruzi* carriers in the long term. By contrast, it was demonstrated that benznidazole monotherapy outperformed posaconazole, with high RT-PCR conversion rates lasting up to a year. However, in 32% of instances, posaconazole side effects resulted in medication termination (72). The various regimens had an acceptable safety profile in the clinical research evaluating fexinidazole; nonetheless, they were ineffective in treating *T. cruzi* infection. This has led to the discontinuation of fexinidazole monotherapy development as a treatment for *T. cruzi* infection (73). These results highlight the necessity of investigating alternative therapeutic approaches to treat CD as well as the significance of conducting thorough assessments of the safety and effectiveness of medicines in clinical trials.

Both benznidazole and nifurtimox have been demonstrated to be more successful in lowering parasitemia during the acute phase of CD. These drugs work by preventing the parasite from synthesizing its DNA, which prevents it from multiplying and spreading throughout the host (74). Furthermore, it has been noted that these medications can cause oxidative damage in the parasite, which helps to eradicate it (75,76). Nevertheless, in the chronic stage of CD, only limited effectiveness of these treatments has been noted; this could be because of the parasite’s persistence in some body tissues, its difficulty entering those tissues, the host’s weakened immune system, and the disease-related tissue damage (77,78). Consequently, the primary focus of current research is on treating the chronic phase of CD, which is not treated by conventional medications. As was previously indicated, VNI has demonstrated encouraging outcomes for CD’s chronic phase. This compound inhibits the Trypanosomatidae enzyme CYP51 (79), which prevents the parasites from synthesizing vital sterols and ultimately compromises the integrity of their cell membranes, killing them (80). It also lessens cardiac fibrosis and inflammation, which may indicate a possibility for stopping or healing heart damage brought on by CD (45). With a wide range of activity against different strains of *Trypanosoma* infections, including resistant ones, and the potential for improved pharmacokinetics and reduced side effects (81–83), this class of inhibitor may offer safer and more efficient alternatives for treating CD during its chronic stage.

Within this species, the strains of *T. cruzi* exhibit unique genetic variations. Regarding their genome, virulence, medication resistance, and capacity to elude the host immune system, each of these strains is distinct (84,85). These variations may have an impact on how CD progresses and how affected individuals respond to treatment (86). The main strains of *T. cruzi* studied in the meta-analysis were Tulahuen and Y, for the chronic and acute phases, respectively. The Y strain, recognized for its high virulence, is preferably employed in models attempting to reproduce the acute phase of the disease, allowing the evaluation of therapy efficacy in decreasing parasitemia and initial infection symptoms (87). On the other hand, research examining the long-term pathogenesis and progression of the disease, as well as the evaluation of therapeutic interventions targeted at lowering parasitic burden and preventing or reversing associated organic damage, utilize the Tulahuen strain, which is recognized for its adaptability and ability to induce a stable chronic infection (88,89).

Conversely, concerning the animal models utilized to assess novel medications against CD, we may bring up the instance of dogs, who serve as significant parasite reservoirs (90). A canine model that accurately mimics human illness has been developed, which makes it easier to assess novel medicinal agents. Routine effectiveness trials are hampered by the high cost and extended lifespan of dogs (91,92). Non-human primates have also been considered possible models, although their usage is restricted due to expense, lack of validation, and ethical concerns (93,94). On the other hand, murine models remain the most popular because of their affordability, portability, and capacity to replicate several facets of human illness (16,95). As demonstrated by the fact that all of the preclinical investigations in this review were carried out in mouse models, they are therefore invaluable resources for researching *T. cruzi* infection and assessing novel antiparasitic medications.

### Limitations and Strengths

The influence on the validity, generalization, and robustness of the results is the intrinsic limitation of the smaller number of scientific publications (96), as is the case in this meta-analysis. A limited data set could not include all available evidence on the topic in question, requiring a cautious interpretation of the results (97). Therefore, it is necessary to support additional research on new treatment options for the CD to overcome this deficiency and increase the available scientific evidence. The current meta-analysis enables a thorough assessment of therapy alternatives in a standardized and controlled setting by concentrating on preclinical studies utilizing animal models, laying a strong platform for further research and therapeutic development. Additionally, complex statistical methods, such as those included in the *"metafor"* package, enable in-depth data analysis, making it easier to control study heterogeneity and accurately estimate treatment effects (98). This helps identify patterns and trends in the effectiveness of treatments, which in turn helps direct decision-making and design of future clinical trials.

### Implications for future research

To identify potential compounds and therapeutic targets, extensive research, including drug discovery approaches, is necessary (99). Similar to the studies included in this meta-analysis, potential therapeutic options should be evaluated in preclinical models, such as cell cultures and animal disease models, to determine their efficacy and safety (100). Subsequently, substances that have potential in preclinical research may proceed to clinical trials. There are several important reasons why many therapeutic approaches that show promise in preclinical research are never tested in clinical trials. Initially, moving from preclinical research to clinical trials is an expensive and complex procedure that requires meticulous preparation and substantial monetary means (101). Clinical studies to evaluate potential drugs can often be hampered by a lack of money or investment from the pharmaceutical industry or funding bodies (102,103). Before starting clinical trials, regulatory and safety issues also need to be resolved. Failure to do so could cause further delays or obstacles in transitioning a promising therapy from preclinical to clinical development (104). Additionally, the availability of resources and motivation to conduct clinical trials may be impacted by the lack of interest or priority of the scientific community or the pharmaceutical industry regarding a particular condition, such as CD (105,106).

## Conclusion

This meta-analysis has demonstrated that novel therapeutic options exist that are successful in treating CD; however, most of them are still in the preclinical development stage, and those that have advanced to the clinical trial stage have not demonstrated the best outcomes, meaning that CD treatment remains unresolved. With the primary goal of helping the CD population, the academic community and pharmaceutical companies must collaborate in creating new drugs, as well as continue and apply research that has produced promising results that could hasten the discovery and availability of more effective treatments.

## Author Contributions

Conceptualization: M.A.C.-P. and M.A.C.-F.; data curation: M.A.C.-P., L.Y.M.-L, B.M.R.-P., and L.D.G.M.; formal analysis: M.A.C.-P. and M.A.C.-F.; funding acquisition: M.A.C.-P., E.A.F.C., and M.A.C.-F.; investigation: L.D.G.M., H.L.B.C, A.S.G, R.A.M.D, R.C.G., and E.A.F.C.; methodology: M.A.C.-P. and M.A.C.-F.; writing—review and editing: L.D.G.M., H.L.B.C, A.S.G, R.A.M.D, R.C.G., and E.A.F.C. All authors have read and agreed to the published version of the manuscript.

## Institutional Review Board Statement

Not applicable.

## Informed Consent Statement

Not applicable.

## Data Availability Statement

Not applicable.

## Acknowledgments

LDGM would like to thank the program PROCIENCIA from CONCYTEC Contrato No PE501084367-2023-PROCIENCIA esquema E067-2023-01 for his fellowship. ASG, RAMA, RCG, and EAFC thank CNPQ for their PQ/DT Fellowship.

## Abbreviations

The following abbreviations are used in this study.

CD: Chagas disease
CI: Confidence interval
c_neg_: Control group negative
c_pos_: Control group positive
DNA: Deoxyribonucleic acid
_I_2: I-squared
INPLASY: International Platform of Registered Systematic Review and Meta-analysis Protocols
MeSH: Medical Subject Headings
PRISMA: Preferred Reporting Items for Systematic Reviews and Meta-Analyses
Q: Q value
RR: Risk Ratio
_T_2: tau-squared
t_neg_: Treated group negative
t_pos_: Treated group positive
*T. cruzi*: *Trypanosoma cruzi*
WHO: World Health Organization

---

**Table S1.**
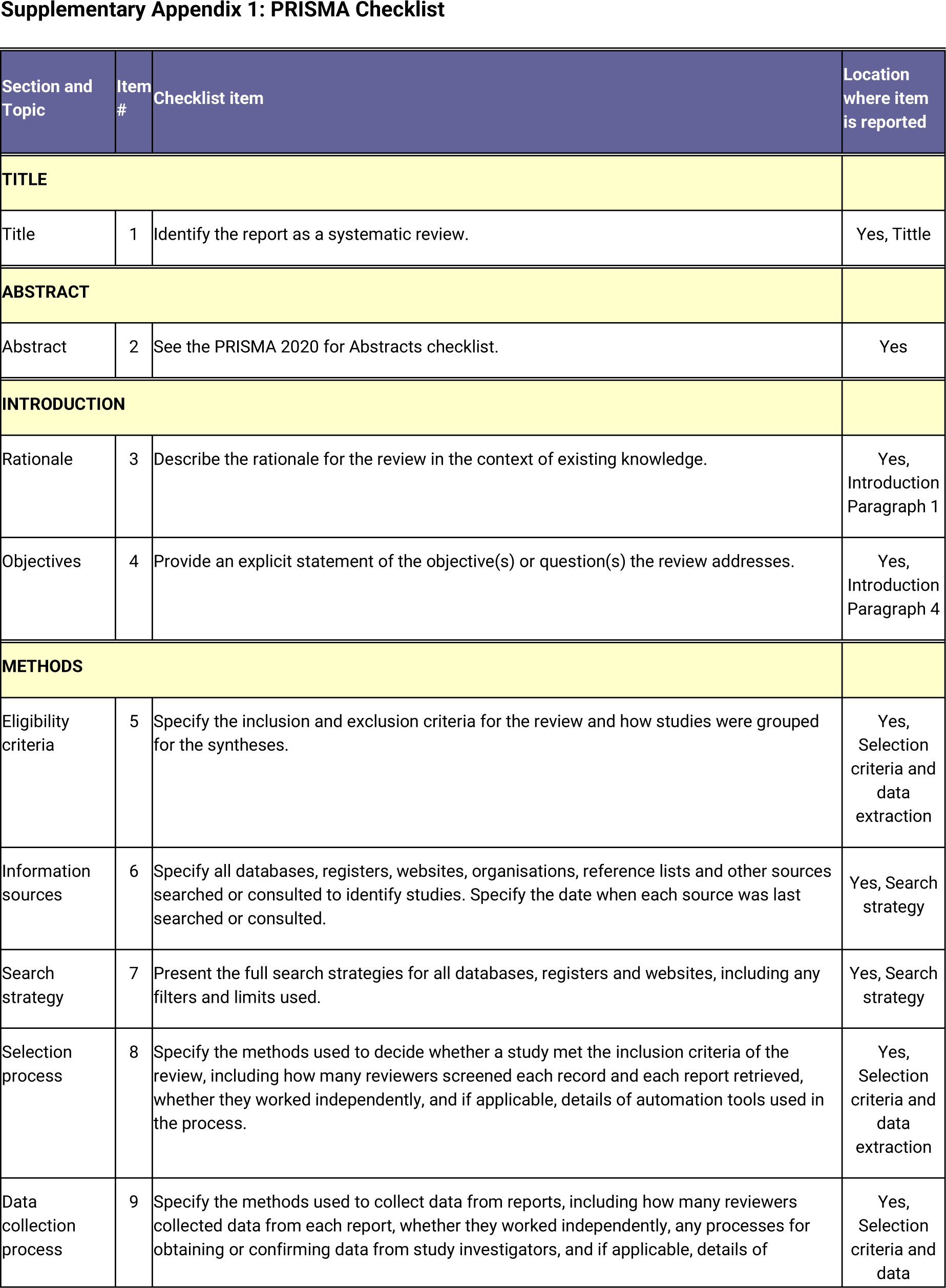
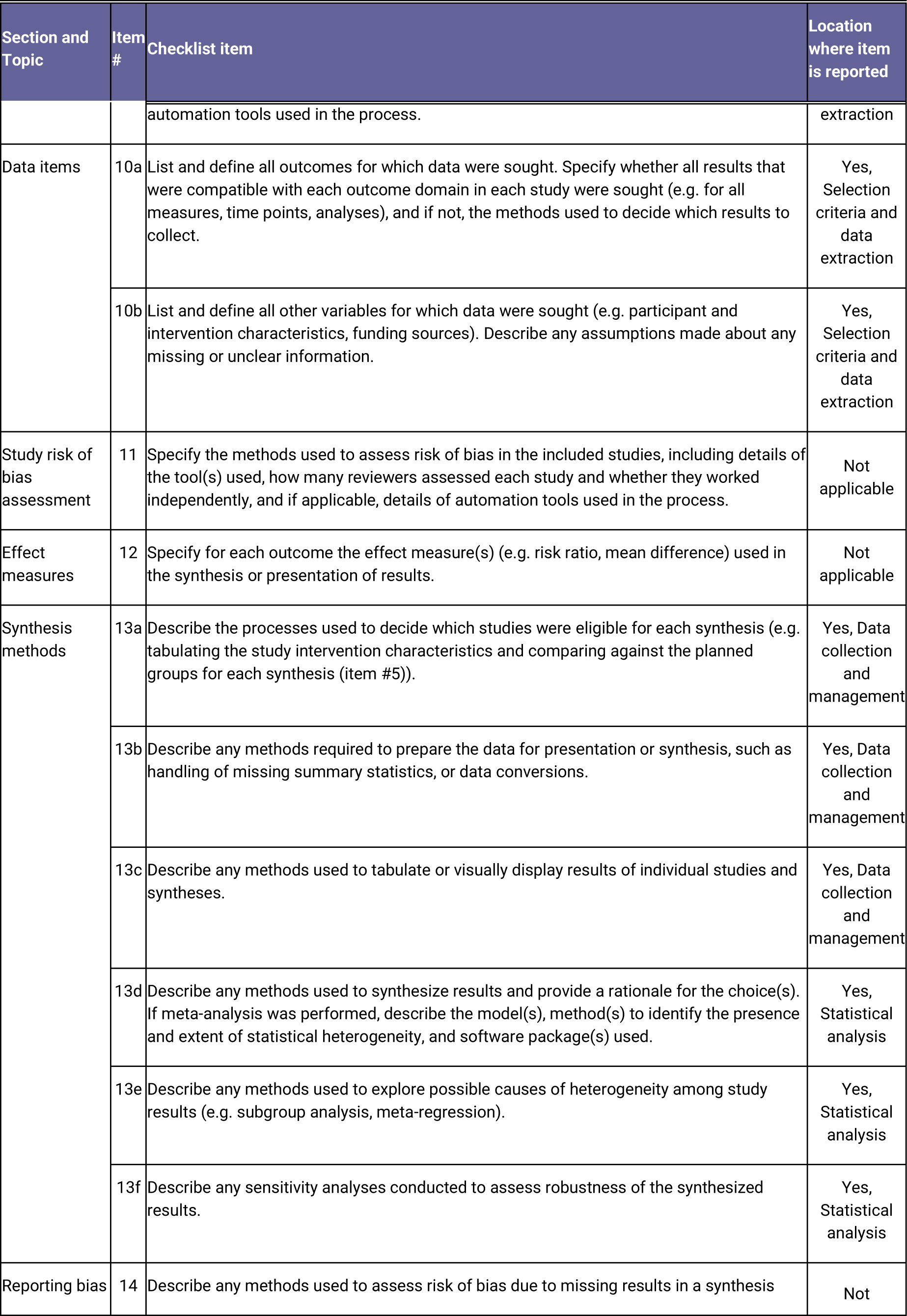
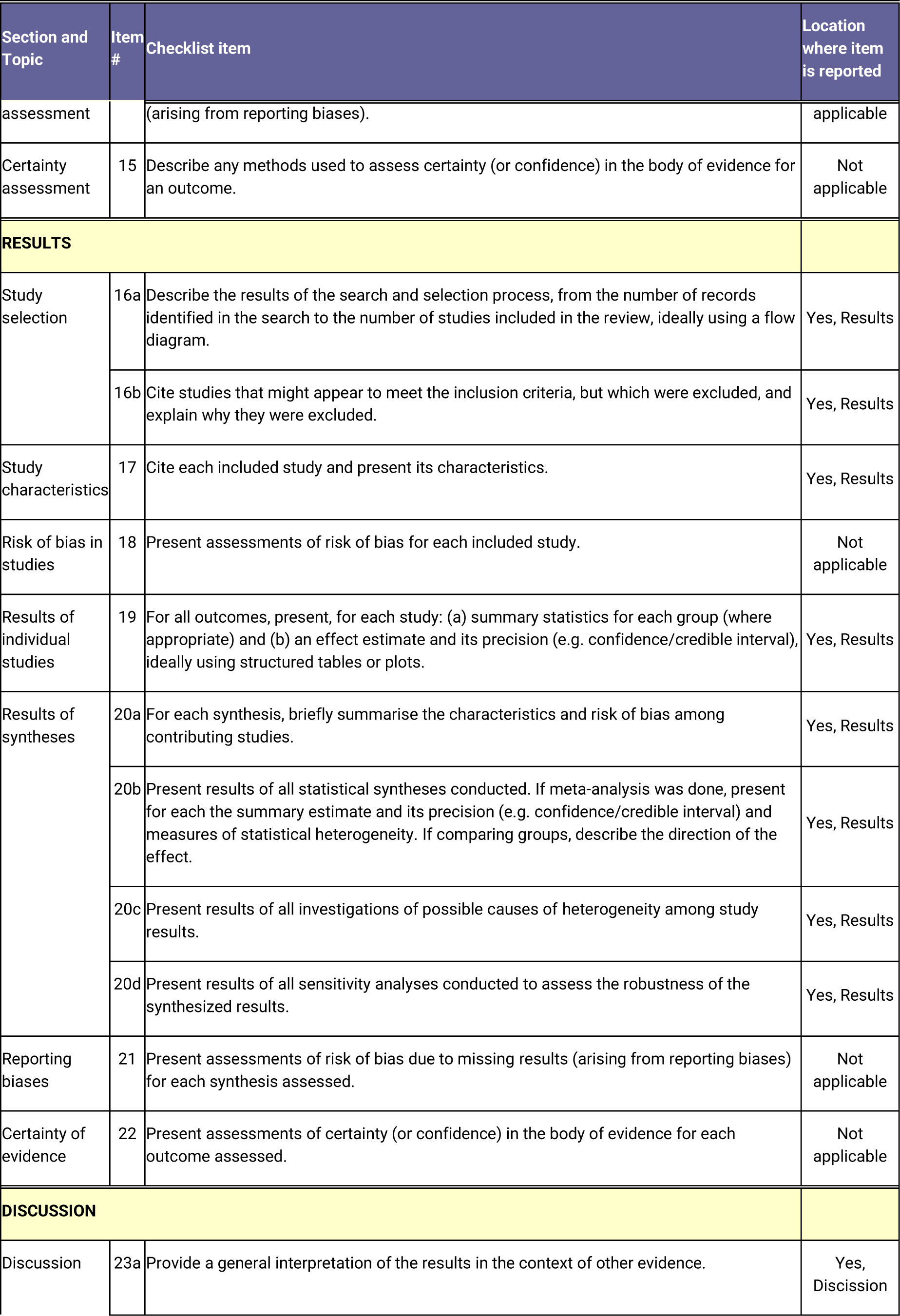
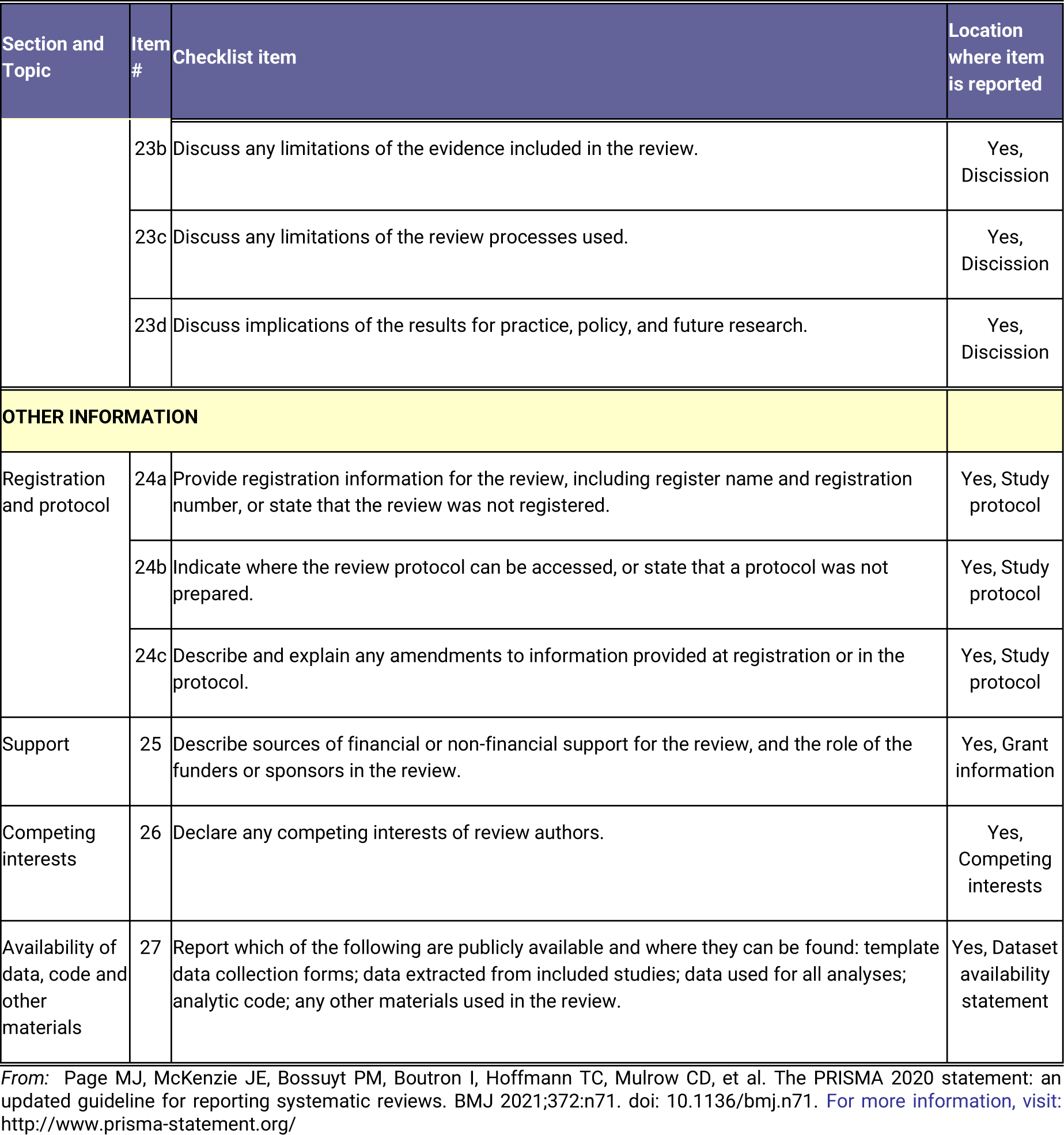
PRISMA 2020 Checklist.

## References

1. Bonney KM. Chagas disease in the 21st Century: a public health success or an emerging threat? Parasite [Internet]. 2014 Mar 10;21(11):1–10. Available from: http://www.parasite-journal.org/10.1051/parasite/2014012

2. Word Health Organization. Chagas’ Disease (American Trypanosomiasis) Fact Sheet [Internet]. 2023 [cited 2024 Feb 23]. Available from: https://www.who.int/news-room/fact-sheets/detail/chagas-disease-(american-trypanosomiasis)

3. Mackey TK, Liang BA, Cuomo R, Hafen R, Brouwer KC, Lee DE. Emerging and Reemerging Neglected Tropical Diseases: a Review of Key Characteristics, Risk Factors, and the Policy and Innovation Environment. Clin Microbiol Rev [Internet]. 2014 Oct;27(4):949–79. Available from: https://journals.asm.org/doi/10.1128/CMR.00045-14

4. Suárez C, Nolder D, García-Mingo A, Moore DA, Chiodini PL. Diagnosis and Clinical Management of Chagas Disease: An Increasing Challenge in Non-Endemic Areas. Res Rep Trop Med [Internet]. 2022 Jul;13:25–40. Available from: https://www.dovepress.com/diagnosis-and-clinical-management-of-chagas-disease-an-increasing-chal-peer-reviewed-fulltext-article-RRTM

5. Echavarría NG, Echeverría LE, Stewart M, Gallego C, Saldarriaga C. Chagas Disease: Chronic Chagas Cardiomyopathy. Curr Probl Cardiol [Internet]. 2021 Mar;46(3):100507. Available from: https://linkinghub.elsevier.com/retrieve/pii/S014628061930180X

6. Iglesias-Rus L, Romay-Barja M, Boquete T, Benito A, Blasco-Hernández T. The role of the first level of health care in the approach to Chagas disease in a non-endemic country. Buekens P, editor. PLoS Negl Trop Dis [Internet]. 2019 Dec 16;13(12):e0007937. Available from: https://dx.plos.org/10.1371/journal.pntd.0007937

7. Henao-Martínez AF, Colborn K, Parra-Henao G. Overcoming research barriers in Chagas disease—designing effective implementation science. Parasitol Res [Internet]. 2017 Jan 22;116(1):35–44. Available from: http://link.springer.com/10.1007/s00436-016-5291-z

8. Alonso-Padilla J, Cortés-Serra N, Pinazo MJ, Bottazzi ME, Abril M, Barreira F, et al. Strategies to enhance access to diagnosis and treatment for Chagas disease patients in Latin America. Expert Rev Anti Infect Ther [Internet]. 2019 Mar 4;17(3):145–57. Available from: https://www.tandfonline.com/doi/full/10.1080/14787210.2019.1577731

9. Vermelho AB, Rodrigues GC, Supuran CT. Why hasn’t there been more progress in new Chagas disease drug discovery? Expert Opin Drug Discov [Internet]. 2020 Feb 1;15(2):145–58. Available from: https://www.tandfonline.com/doi/full/10.1080/17460441.2020.1681394

10. Ribeiro V, Dias N, Paiva T, Hagström-Bex L, Nitz N, Pratesi R, et al. Current trends in the pharmacological management of Chagas disease. Int J Parasitol Drugs Drug Resist [Internet]. 2020 Apr;12:7–17. Available from: https://linkinghub.elsevier.com/retrieve/pii/S2211320719301447

11. Bermudez J, Davies C, Simonazzi A, Pablo Real J, Palma S. Current drug therapy and pharmaceutical challenges for Chagas disease. Acta Trop [Internet]. 2016 Apr;156:1–16. Available from: https://linkinghub.elsevier.com/retrieve/pii/S0001706X15301947

12. Donadeu M, Nwankpa N, Abela-Ridder B, Dungu B. Strategies to increase adoption of animal vaccines by smallholder farmers with focus on neglected diseases and marginalized populations. Rupprecht CE, editor. PLoS Negl Trop Dis [Internet]. 2019 Feb 7;13(2):e0006989. Available from: https://dx.plos.org/10.1371/journal.pntd.0006989

13. Acevedo GR, Girard MC, Gómez KA. The Unsolved Jigsaw Puzzle of the Immune Response in Chagas Disease. Front Immunol [Internet]. 2018 Aug 24;9. Available from: https://www.frontiersin.org/article/10.3389/fimmu.2018.01929/full

14. Weng H-B, Chen H-X, Wang M-W. Innovation in neglected tropical disease drug discovery and development. Infect Dis Poverty [Internet]. 2018 Dec 18;7(1):67. Available from: https://idpjournal.biomedcentral.com/articles/10.1186/s40249-018-0444-1

15. Echeverría LE, Marcus R, Novick G, Sosa-Estani S, Ralston K, Zaidel EJ, et al. WHF IASC Roadmap on Chagas Disease. Glob Heart [Internet]. 2020 Mar 30;15(1):26. Available from: https://globalheartjournal.com/article/10.5334/gh.484/

16. Chatelain E, Scandale I. Animal models of Chagas disease and their translational value to drug development. Expert Opin Drug Discov [Internet]. 2020 Dec 1;15(12):1381–402. Available from: https://www.tandfonline.com/doi/full/10.1080/17460441.2020.1806233

17. Martínez-Peinado N, Cortes-Serra N, Losada-Galvan I, Alonso-Vega C, Urbina JA, Rodríguez A, et al. Emerging agents for the treatment of Chagas disease: what is in the preclinical and clinical development pipeline? Expert Opin Investig Drugs [Internet]. 2020 Sep 1;29(9):947–59. Available from: https://www.tandfonline.com/doi/full/10.1080/13543784.2020.1793955

18. García-Huertas P, Cardona-Castro N. Advances in the treatment of Chagas disease: Promising new drugs, plants and targets. Biomed Pharmacother [Internet]. 2021 Oct;142:112020. Available from: https://linkinghub.elsevier.com/retrieve/pii/S0753332221008039

19. Page MJ, Moher D, Bossuyt PM, Boutron I, Hoffmann TC, Mulrow CD, et al. PRISMA 2020 explanation and elaboration: updated guidance and exemplars for reporting systematic reviews. BMJ [Internet]. 2021 Mar 29;372:n160. Available from: https://www.bmj.com/lookup/doi/10.1136/bmj.n160

20. van Eck NJ, Waltman L. Citation-based clustering of publications using CitNetExplorer and VOSviewer. Scientometrics [Internet]. 2017 May 27;111(2):1053–70. Available from: http://link.springer.com/10.1007/s11192-017-2300-7

21. National Library of Medicine. Treatment Outcome [Internet]. MeSH Terms. 1992 [cited 2024 Feb 28]. Available from: https://www.ncbi.nlm.nih.gov/mesh/?term=treatment+outcome

22. Drevon D, Fursa SR, Malcolm AL. Intercoder Reliability and Validity of WebPlotDigitizer in Extracting Graphed Data. Behav Modif [Internet]. 2017 Mar 22;41(2):323–39. Available from: http://journals.sagepub.com/doi/10.1177/0145445516673998

23. Rohatgi A. WebPlotDigitizer [Internet]. California, USA; 2024. Available from: https://automeris.io/WebPlotDigitizer

24. Rhim HC, Kim SJ, Park J, Jang K-M. Effect of citrulline on post-exercise rating of perceived exertion, muscle soreness, and blood lactate levels: A systematic review and meta-analysis. J Sport Heal Sci [Internet]. 2020 Dec;9(6):553–61. Available from: https://linkinghub.elsevier.com/retrieve/pii/S2095254620300168

25. Vaka R, Remortel S Van, Ly V, Davis DR. Extracellular vesicle therapy for non-ischemic heart failure: A systematic review of preclinical studies. Extracell Vesicle [Internet]. 2022 Dec;1:100009. Available from: https://linkinghub.elsevier.com/retrieve/pii/S277304172200004X

26. Reis DJ, Ilardi SS, Punt SEW. The anxiolytic effect of probiotics: A systematic review and meta-analysis of the clinical and preclinical literature. Foster J, editor. PLoS One [Internet]. 2018 Jun 20;13(6):e0199041. Available from: https://dx.plos.org/10.1371/journal.pone.0199041

27. Viechtbauer W. Conducting Meta-Analyses in R with the metafor Package. J Stat Softw [Internet]. 2010;36(3):1–48. Available from: http://www.jstatsoft.org/v36/i03/

28. Polanin JR, Hennessy EA, Tanner-Smith EE. A Review of Meta-Analysis Packages in R. J Educ Behav Stat [Internet]. 2017 Apr 30;42(2):206–42. Available from: http://journals.sagepub.com/doi/10.3102/1076998616674315

29. Borenstein M, Hedges L V., Higgins JPT, Rothstein HR. A basic introduction to fixed-effect and random-effects models for meta-analysis. Res Synth Methods [Internet]. 2010 Apr;1(2):97–111. Available from: https://onlinelibrary.wiley.com/doi/10.1002/jrsm.12

30. Ruppar T. Meta-analysis: How to quantify and explain heterogeneity? Eur J Cardiovasc Nurs [Internet]. 2020 Oct 5;19(7):646–52. Available from: https://academic.oup.com/eurjcn/article/19/7/646-652/5950370

31. von Hippel PT. The heterogeneity statistic I2 can be biased in small meta-analyses. BMC Med Res Methodol [Internet]. 2015 Dec 14;15(1):35. Available from: https://bmcmedresmethodol.biomedcentral.com/articles/10.1186/s12874-015-0024-z

32. Böhning D, Lerdsuwansri R, Holling H. Some general points on the I2-measure of heterogeneity in meta-analysis. Metrika [Internet]. 2017 Nov 22;80(6–8):685–95. Available from: http://link.springer.com/10.1007/s00184-017-0622-3

33. Borenstein M. Avoiding common mistakes in meta-analysis: Understanding the distinct roles of Q, I-squared, tau-squared, and the prediction interval in reporting heterogeneity. Res Synth Methods [Internet]. 2023 Nov 8; Available from: https://onlinelibrary.wiley.com/doi/10.1002/jrsm.1678

34. Vesterinen HM, Sena ES, Egan KJ, Hirst TC, Churolov L, Currie GL, et al. Meta-analysis of data from animal studies: A practical guide. J Neurosci Methods [Internet]. 2014 Jan;221:92–102. Available from: https://linkinghub.elsevier.com/retrieve/pii/S016502701300321X

35. D’Arrigo G, Gori M, Pitino A, Tsalikakis DG, Liakopoulos V, Roumeliotis S, et al. Measures of frequency and effect in clinical research. Int Urol Nephrol [Internet]. 2023 May 10;55(12):3147–52. Available from: https://link.springer.com/10.1007/s11255-023-03626-w

36. Andrade C. Understanding Relative Risk, Odds Ratio, and Related Terms: As Simple as It Can Get. J Clin Psychiatry [Internet]. 2015 Jul 22;76(07):e857–61. Available from: https://www.psychiatrist.com/jcp/understanding-relative-risk-odds-ratio-related-terms

37. Cencig S, Coltel N, Truyens C, Carlier Y. Evaluation of benznidazole treatment combined with nifurtimox, posaconazole or AmBisome® in mice infected with Trypanosoma cruzi strains. Int J Antimicrob Agents [Internet]. 2012 Dec;40(6):527–32. Available from: https://linkinghub.elsevier.com/retrieve/pii/S0924857912003366

38. Diniz L de F, Urbina JA, de Andrade IM, Mazzeti AL, Martins TAF, Caldas IS, et al. Benznidazole and Posaconazole in Experimental Chagas Disease: Positive Interaction in Concomitant and Sequential Treatments. Rodrigues MM, editor. PLoS Negl Trop Dis [Internet]. 2013 Aug 15;7(8):e2367. Available from: https://dx.plos.org/10.1371/journal.pntd.0002367

39. Molina J. Cure of experimental Chagas’ disease by the bis-triazole DO870 incorporated into “stealth” polyethyleneglycol-polylactide nanospheres. J Antimicrob Chemother [Internet]. 2001 Jan 1;47(1):101–4. Available from: https://academic.oup.com/jac/article-lookup/doi/10.1093/jac/47.1.101

40. Ndao M, Beaulieu C, Black WC, Isabel E, Vasquez-Camargo F, Nath-Chowdhury M, et al. Reversible Cysteine Protease Inhibitors Show Promise for a Chagas Disease Cure. Antimicrob Agents Chemother [Internet]. 2014 Feb;58(2):1167–78. Available from: https://journals.asm.org/doi/10.1128/AAC.01855-13

41. Matsuo AL, Silva LS, Torrecilhas AC, Pascoalino BS, Ramos TC, Rodrigues EG, et al. In Vitro and In Vivo Trypanocidal Effects of the Cyclopalladated Compound 7a, a Drug Candidate for Treatment of Chagas’ Disease. Antimicrob Agents Chemother [Internet]. 2010 Aug;54(8):3318–25. Available from: https://journals.asm.org/doi/10.1128/AAC.00323-10

42. Bahia MT, Andrade IM de, Martins TAF, Nascimento ÁF da S do, Diniz L de F, Caldas IS, et al. Fexinidazole: A Potential New Drug Candidate for Chagas Disease. Pollastri MP, editor. PLoS Negl Trop Dis [Internet]. 2012 Nov 1;6(11):e1870. Available from: https://dx.plos.org/10.1371/journal.pntd.0001870

43. da Silva CF, Batista D da GJ, De Araújo JS, Batista MM, Lionel J, de Souza EM, et al. Activities of Psilostachyin A and Cynaropicrin against Trypanosoma cruzi In Vitro and In Vivo. Antimicrob Agents Chemother [Internet]. 2013 Nov;57(11):5307–14. Available from: https://journals.asm.org/doi/10.1128/AAC.00595-13

44. Santos CD, Loria RM, Rodrigues Oliveira LG, Collins Kuehn C, Alonso Toldo MP, Albuquerque S, et al. Effects of dehydroepiandrosterone-sulfate (DHEA-S) and benznidazole treatments during acute infection of two different Trypanosoma cruzi strains. Immunobiology [Internet]. 2010 Dec;215(12):980–6. Available from: https://linkinghub.elsevier.com/retrieve/pii/S0171298509001727

45. Villalta F, Dobish MC, Nde PN, Kleshchenko YY, Hargrove TY, Johnson CA, et al. VNI Cures Acute and Chronic Experimental Chagas Disease. J Infect Dis [Internet]. 2013 Aug 1;208(3):504–11. Available from: https://academic.oup.com/jid/article-lookup/doi/10.1093/infdis/jit042

46. Saraiva J, Lira AAM, Esperandim VR, da Silva Ferreira D, Ferraudo AS, Bastos JK, et al. (−)−Hinokinin-loaded poly(d,l-lactide-co-glycolide) microparticles for Chagas disease. Parasitol Res [Internet]. 2010 Feb 28;106(3):703–8. Available from: http://link.springer.com/10.1007/s00436-010-1725-1

47. Gobbi P, Baez A, Lo Presti MS, Fernández AR, Enders JE, Fretes R, et al. Association of clomipramine and allopurinol for the treatment of the experimental infection with Trypanosoma cruzi. Parasitol Res [Internet]. 2010 Oct 3;107(5):1279–83. Available from: http://link.springer.com/10.1007/s00436-010-2002-z

48. de Oliveira FL, Araújo-Jorge TC, de Souza EM, de Oliveira GM, Degrave WM, Feige J-J, et al. Oral Administration of GW788388, an Inhibitor of Transforming Growth Factor Beta Signaling, Prevents Heart Fibrosis in Chagas Disease. Rodrigues MM, editor. PLoS Negl Trop Dis [Internet]. 2012 Jun 12;6(6):e1696. Available from: https://dx.plos.org/10.1371/journal.pntd.0001696

49. Dumonteil E, Escobedo-Ortegon J, Reyes-Rodriguez N, Arjona-Torres A, Ramirez-Sierra MJ. Immunotherapy of Trypanosoma cruzi Infection with DNA Vaccines in Mice. Infect Immun [Internet]. 2004 Jan;72(1):46–53. Available from: https://journals.asm.org/doi/10.1128/IAI.72.1.46-53.2004

50. Mattia G, Puglisi R, Ascione B, Malorni W, Carè A, Matarrese P. Cell death-based treatments of melanoma: conventional treatments and new therapeutic strategies. Cell Death Dis [Internet]. 2018 Jan 25;9(2):112. Available from: https://www.nature.com/articles/s41419-017-0059-7

51. Wang W, Peng D, Baptista RP, Li Y, Kissinger JC, Tarleton RL. Strain-specific genome evolution in Trypanosoma cruzi, the agent of Chagas disease. Siegel N, editor. PLOS Pathog [Internet]. 2021 Jan 28;17(1):e1009254. Available from: https://dx.plos.org/10.1371/journal.ppat.1009254

52. Altamura F, Rajesh R, Catta-Preta CMC, Moretti NS, Cestari I. The current drug discovery landscape for trypanosomiasis and leishmaniasis: Challenges and strategies to identify drug targets. Drug Dev Res [Internet]. 2022 Apr 6;83(2):225–52. Available from: https://onlinelibrary.wiley.com/doi/10.1002/ddr.21664

53. Zingales B. Trypanosoma cruzi genetic diversity: Something new for something known about Chagas disease manifestations, serodiagnosis and drug sensitivity. Acta Trop [Internet]. 2018 Aug;184:38–52. Available from: https://linkinghub.elsevier.com/retrieve/pii/S0001706X17304266

54. Manne JM, Snively CS, Ramsey JM, Salgado MO, Bärnighausen T, Reich MR. Barriers to Treatment Access for Chagas Disease in Mexico. Franco-Paredes C, editor. PLoS Negl Trop Dis [Internet]. 2013 Oct 17;7(10):e2488. Available from: https://dx.plos.org/10.1371/journal.pntd.0002488

55. Wirtz VJ, Manne-Goehler J, Reich MR. Access to Care for Chagas Disease in the United States: A Health Systems Analysis. Am J Trop Med Hyg [Internet]. 2015 Jul 8;93(1):108–13. Available from: https://ajtmh.org/doi/10.4269/ajtmh.14-0826

56. Mills RM. Chagas Disease: Epidemiology and Barriers to Treatment. Am J Med [Internet]. 2020 Nov;133(11):1262–5. Available from: https://linkinghub.elsevier.com/retrieve/pii/S0002934320305209

57. Leisinger KM. Poverty, Disease, and Medicines in Low- and Middle-Income Countries. Bus Prof Ethics J [Internet]. 2012;31(1):135–85. Available from: http://www.pdcnet.org/oom/service?url_ver=Z39.88-2004&rft_val_fmt=&rft.imuse_id=bpej_2012_0031_0001_0135_0185&svc_id=info:www.pdcnet.org/collection

58. Olliaro PL, Kuesel AC, Reeder JC. A Changing Model for Developing Health Products for Poverty-Related Infectious Diseases. Geary TG, editor. PLoS Negl Trop Dis [Internet]. 2015 Jan 8;9(1):e3379. Available from: https://dx.plos.org/10.1371/journal.pntd.0003379

59. Martín-Escolano J, Medina-Carmona E, Martín-Escolano R. Chagas Disease: Current View of an Ancient and Global Chemotherapy Challenge. ACS Infect Dis [Internet]. 2020 Nov 13;6(11):2830–43. Available from: https://pubs.acs.org/doi/10.1021/acsinfecdis.0c00353

60. Chatelain E. Chagas Disease Drug Discovery: Toward a New Era. SLAS Discov [Internet]. 2015 Jan;20(1):22–35. Available from: https://linkinghub.elsevier.com/retrieve/pii/S2472555222071738

61. Muñoz-Calderón A, Ramírez JL, Díaz-Bello Z, Alarcón de Noya B, Noya O, Schijman AG. Genetic Characterization of Trypanosoma cruzi I Populations from an Oral Chagas Disease Outbreak in Venezuela: Natural Resistance to Nitroheterocyclic Drugs. ACS Infect Dis [Internet]. 2023 Mar 10;9(3):582–92. Available from: https://pubs.acs.org/doi/10.1021/acsinfecdis.2c00569

62. Fonseca B de P, Albuquerque PC, Zicker F. Neglected tropical diseases in Brazil: lack of correlation between disease burden, research funding and output. Trop Med Int Heal [Internet]. 2020 Nov 17;25(11):1373–84. Available from: https://www.intechopen.com/online-first/1155921

63. Schofield CJ, Jannin J, Salvatella R. The future of Chagas disease control. Trends Parasitol [Internet]. 2006 Dec;22(12):583–8. Available from: https://linkinghub.elsevier.com/retrieve/pii/S1471492206002601

64. de Carvalho ACC, de Souza W. The evolution of Brazilian Health Sciences and the present situation. Lancet Reg Heal - Am [Internet]. 2021 Nov;3:100044. Available from: https://linkinghub.elsevier.com/retrieve/pii/S2667193X21000363

65. Moncayo Á, Silveira AC. Current epidemiological trends of Chagas disease in Latin America and future challenges. In: American Trypanosomiasis Chagas Disease [Internet]. Elsevier; 2017. p. 59–88. Available from: https://linkinghub.elsevier.com/retrieve/pii/B9780128010297000046

66. Ng-Kamstra JS, Greenberg SLM, Abdullah F, Amado V, Anderson GA, Cossa M, et al. Global Surgery 2030: a roadmap for high income country actors. BMJ Glob Heal [Internet]. 2016 Apr;1(1):e000011. Available from: https://gh.bmj.com/lookup/doi/10.1136/bmjgh-2015-000011

67. Bern C, Messenger LA, Whitman JD, Maguire JH. Chagas Disease in the United States: a Public Health Approach. Clin Microbiol Rev [Internet]. 2019 Dec 18;33(1). Available from: https://journals.asm.org/doi/10.1128/CMR.00023-19

68. Bern C, Montgomery SP, Herwaldt BL, Rassi A, Marin-Neto JA, Dantas RO, et al. Evaluation and Treatment of Chagas Disease in the United States. JAMA [Internet]. 2007 Nov 14;298(18):2171. Available from: http://jama.jamanetwork.com/article.aspx?doi=10.1001/jama.298.18.2171

69. Gascon J, Bern C, Pinazo M-J. Chagas disease in Spain, the United States and other non-endemic countries. Acta Trop [Internet]. 2010 Jul;115(1–2):22–7. Available from: https://linkinghub.elsevier.com/retrieve/pii/S0001706X09001995

70. Pérez-Molina JA, Norman F, López-Vélez R. Chagas Disease in Non-Endemic Countries: Epidemiology, Clinical Presentation and Treatment. Curr Infect Dis Rep [Internet]. 2012 Jun 3;14(3):263–74. Available from: http://link.springer.com/10.1007/s11908-012-0259-3

71. Castillo-Riquelme M. Chagas disease in non-endemic countries. Lancet Glob Heal [Internet]. 2017 Apr;5(4):e379–80. Available from: https://linkinghub.elsevier.com/retrieve/pii/S2214109X17300906

72. Morillo CA, Waskin H, Sosa-Estani S, del Carmen Bangher M, Cuneo C, Milesi R, et al. Benznidazole and Posaconazole in Eliminating Parasites in Asymptomatic T. Cruzi Carriers: The STOP-CHAGAS Trial. J Am Coll Cardiol [Internet]. 2017 Feb;69(8):939–47. Available from: https://linkinghub.elsevier.com/retrieve/pii/S0735109717301158

73. Pinazo M-J, Forsyth C, Losada I, Esteban ET, García-Rodríguez M, Villegas ML, et al. Efficacy and safety of fexinidazole for treatment of chronic indeterminate Chagas disease (FEXI-12): a multicentre, randomised, double-blind, phase 2 trial. Lancet Infect Dis [Internet]. 2024 Jan; Available from: https://linkinghub.elsevier.com/retrieve/pii/S1473309923006515

74. Varikuti S, Jha BK, Volpedo G, Ryan NM, Halsey G, Hamza OM, et al. Host-Directed Drug Therapies for Neglected Tropical Diseases Caused by Protozoan Parasites. Front Microbiol [Internet]. 2018 Nov 30;9. Available from: https://www.frontiersin.org/article/10.3389/fmicb.2018.02655/full

75. Boiani M, Piacenza L, Hernández P, Boiani L, Cerecetto H, González M, et al. Mode of action of Nifurtimox and N-oxide-containing heterocycles against Trypanosoma cruzi: Is oxidative stress involved? Biochem Pharmacol [Internet]. 2010 Jun;79(12):1736–45. Available from: https://linkinghub.elsevier.com/retrieve/pii/S0006295210001036

76. Faundez M, Pino L, Letelier P, Ortiz C, López R, Seguel C, et al. Buthionine Sulfoximine Increases the Toxicity of Nifurtimox and Benznidazole to Trypanosoma cruzi. Antimicrob Agents Chemother [Internet]. 2005 Jan;49(1):126–30. Available from: https://journals.asm.org/doi/10.1128/AAC.49.1.126-130.2005

77. de Oliveira RB, Ballart C, Abràs A, Gállego M, Marin-Neto JA. Chagas Disease: An Unknown and Neglected Disease. In: Chagas Disease [Internet]. Cham: Springer International Publishing; 2020. p. 1–26. Available from: http://link.springer.com/10.1007/978-3-030-44054-1_1

78. López-Vélez R, Norman FF, Bern C. American Trypanosomiasis (Chagas Disease). In: Hunter’s Tropical Medicine and Emerging Infectious Diseases [Internet]. Elsevier; 2020. p. 762–75. Available from: https://linkinghub.elsevier.com/retrieve/pii/B9780323555128001034

79. Lepesheva GI, Villalta F, Waterman MR. Targeting Trypanosoma cruzi Sterol 14α- Demethylase (CYP51). In 2011. p. 65–87. Available from: https://linkinghub.elsevier.com/retrieve/pii/B9780123858634000046

80. de Morais CGV, Castro Lima AK, Terra R, dos Santos RF, Da-Silva SAG, Dutra PML. The Dialogue of the Host-Parasite Relationship: Leishmania spp. and Trypanosoma cruzi Infection. Biomed Res Int [Internet]. 2015;2015:1–19. Available from: http://www.hindawi.com/journals/bmri/2015/324915/

81. Soeiro M de NC, de Souza EM, da Silva CF, Batista D da GJ, Batista MM, Pavão BP, et al. In Vitro and In Vivo Studies of the Antiparasitic Activity of Sterol 14α-Demethylase (CYP51) Inhibitor VNI against Drug-Resistant Strains of Trypanosoma cruzi. Antimicrob Agents Chemother [Internet]. 2013 Sep;57(9):4151–63. Available from: https://journals.asm.org/doi/10.1128/AAC.00070-13

82. Guedes-da-Silva FH, Batista DGJ, Da Silva CF, De Araújo JS, Pavão BP, Simões-Silva MR, et al. Antitrypanosomal Activity of Sterol 14α-Demethylase (CYP51) Inhibitors VNI and VFV in the Swiss Mouse Models of Chagas Disease Induced by the Trypanosoma cruzi Y Strain. Antimicrob Agents Chemother [Internet]. 2017 Apr;61(4). Available from: https://journals.asm.org/doi/10.1128/AAC.02098-16

83. Friggeri L, Hargrove TY, Rachakonda G, Blobaum AL, Fisher P, de Oliveira GM, et al. Sterol 14α-Demethylase Structure-Based Optimization of Drug Candidates for Human Infections with the Protozoan Trypanosomatidae. J Med Chem [Internet]. 2018 Dec 13;61(23):10910–21. Available from: https://pubs.acs.org/doi/10.1021/acs.jmedchem.8b01671

84. Junqueira C, Caetano B, Bartholomeu DC, Melo MB, Ropert C, Rodrigues MM, et al. The endless race between Trypanosoma cruzi and host immunity: lessons for and beyond Chagas disease. Expert Rev Mol Med [Internet]. 2010 Sep 15;12:e29. Available from: https://www.cambridge.org/core/product/identifier/S1462399410001560/type/journal_article

85. Jiménez P, Jaimes J, Poveda C, Ramírez JD. A systematic review of the Trypanosoma cruzi genetic heterogeneity, host immune response and genetic factors as plausible drivers of chronic chagasic cardiomyopathy. Parasitology [Internet]. 2019 Mar 13;146(3):269–83. Available from: https://www.cambridge.org/core/product/identifier/S0031182018001506/type/journal_article

86. Santi-Rocca J, Fernandez-Cortes F, Chillón-Marinas C, González-Rubio M-L, Martin D, Gironès N, et al. A multi-parametric analysis of Trypanosoma cruzi infection: common pathophysiologic patterns beyond extreme heterogeneity of host responses. Sci Rep [Internet]. 2017 Aug 21;7(1):8893. Available from: https://www.nature.com/articles/s41598-017-08086-8

87. Mateus J, Guerrero P, Lasso P, Cuervo C, González JM, Puerta CJ, et al. An Animal Model of Acute and Chronic Chagas Disease With the Reticulotropic Y Strain of Trypanosoma cruzi That Depicts the Multifunctionality and Dysfunctionality of T Cells. Front Immunol [Internet]. 2019 Apr 26;10. Available from: https://www.frontiersin.org/article/10.3389/fimmu.2019.00918/full

88. Scarim CB, Jornada DH, Chelucci RC, de Almeida L, dos Santos JL, Chung MC. Current advances in drug discovery for Chagas disease. Eur J Med Chem [Internet]. 2018 Jul;155:824–38. Available from: https://linkinghub.elsevier.com/retrieve/pii/S0223523418305312

89. Bustamante JM, Presti MS Lo, Rivarola HW, Fernández AR, Enders JE, Fretes RE, et al. Treatment with benznidazole or thioridazine in the chronic phase of experimental Chagas disease improves cardiopathy. Int J Antimicrob Agents [Internet]. 2007 Jun 9;29(6):733–7. Available from: https://www.cambridge.org/core/product/identifier/S0031182012001771/type/journal_article

90. Gürtler RE, Cardinal MV. Reservoir host competence and the role of domestic and commensal hosts in the transmission of Trypanosoma cruzi. Acta Trop [Internet]. 2015 Nov;151:32–50. Available from: https://linkinghub.elsevier.com/retrieve/pii/S0001706X1530022X

91. Guedes PM da M, Veloso VM, Tafuri WL, Galvão LM da C, Carneiro CM, Lana M de, et al. The dog as model for chemotherapy of the Chagas’ disease. Acta Trop [Internet]. 2002 Oct;84(1):9–17. Available from: https://linkinghub.elsevier.com/retrieve/pii/S0001706X02001390

92. Rodríguez-Morales O, Roldán F-J, Vargas-Barrón J, Parra-Benítez E, Medina-García M de L, Vergara-Bello E, et al. Echocardiographic Findings in Canine Model of Chagas Disease Immunized with DNA Trypanosoma cruzi Genes. Animals [Internet]. 2020 Apr 9;10(4):648. Available from: https://www.mdpi.com/2076-2615/10/4/648

93. Pung OJ, Hulsebos LH, Kuhn RE. Experimental Chagas’ disease (Trypanosoma cruzi) in the Brazilian squirrel monkey (Saimiri sciureus): Hematology, cardiology, cellular and humoral immune responses. Int J Parasitol [Internet]. 1988 Feb;18(1):115–20. Available from: https://linkinghub.elsevier.com/retrieve/pii/0020751988900458

94. Seah SKK, Marsden PD, Voller A, Pettitt LE. Experimental Trypanosoma cruzi infection in rhesus monkeys—The acute phase. Trans R Soc Trop Med Hyg [Internet]. 1974 Jan;68(1):63–9. Available from: https://academic.oup.com/trstmh/article-lookup/doi/10.1016/0035-9203(74)90254-5

95. Costa SCG da. Mouse as a model for Chagas disease: does mouse represent a good model for Chagas disease? Mem Inst Oswaldo Cruz [Internet]. 1999 Sep;94(suppl 1):269–72. Available from: http://www.scielo.br/scielo.php?script=sci_arttext&pid=S0074-02761999000700045&lng=en&tlng=en

96. Gurevitch J, Koricheva J, Nakagawa S, Stewart G. Meta-analysis and the science of research synthesis. Nature [Internet]. 2018 Mar 8;555(7695):175–82. Available from: https://www.nature.com/articles/nature25753

97. Garg AX, Hackam D, Tonelli M. Systematic Review and Meta-analysis: When One Study Is Just not Enough. Clin J Am Soc Nephrol [Internet]. 2008 Jan;3(1):253–60. Available from: https://journals.lww.com/01277230-200801000-00038

98. Wang N. Conducting Meta-analyses of Proportions in R. J Behav Data Sci [Internet]. 2023;3(2). Available from: https://jbds.isdsa.org/jbds/article/view/60

99. Kiriiri GK, Njogu PM, Mwangi AN. Exploring different approaches to improve the success of drug discovery and development projects: a review. Futur J Pharm Sci [Internet]. 2020 Dec 23;6(1):27. Available from: https://fjps.springeropen.com/articles/10.1186/s43094-020-00047-9

100. Wang H, Brown PC, Chow ECY, Ewart L, Ferguson SS, Fitzpatrick S, et al. 3D cell culture models: Drug pharmacokinetics, safety assessment, and regulatory consideration. Clin Transl Sci [Internet]. 2021 Sep 16;14(5):1659–80. Available from: https://ascpt.onlinelibrary.wiley.com/doi/10.1111/cts.13066

101. Liakos A, Pagkalidou E, Karagiannis T, Malandris K, Avgerinos I, Gigi E, et al. A Simple Guide to Randomized Controlled Trials. Int J Low Extrem Wounds [Internet]. 2024 Feb 28; Available from: http://journals.sagepub.com/doi/10.1177/15347346241236385

102. Mahlich J, Bartol A, Dheban S. Can adaptive clinical trials help to solve the productivity crisis of the pharmaceutical industry? - a scenario analysis. Health Econ Rev [Internet]. 2021 Dec 16;11(1):4. Available from: https://healtheconomicsreview.biomedcentral.com/articles/10.1186/s13561-021-00302-6

103. Schlander M, Hernandez-Villafuerte K, Cheng C-Y, Mestre-Ferrandiz J, Baumann M. How Much Does It Cost to Research and Develop a New Drug? A Systematic Review and Assessment. Pharmacoeconomics [Internet]. 2021 Nov 9;39(11):1243–69. Available from: https://link.springer.com/10.1007/s40273-021-01065-y

104. Darrow JJ, Avorn J, Kesselheim AS. FDA Approval and Regulation of Pharmaceuticals, 1983-2018. JAMA [Internet]. 2020 Jan 14;323(2):164. Available from: https://jamanetwork.com/journals/jama/fullarticle/2758605

105. Jimeno I, Mendoza N, Zapana F, de la Torre L, Torrico F, Lozano D, et al. Social determinants in the access to health care for Chagas disease: A qualitative research on family life in the “Valle Alto” of Cochabamba, Bolivia. Herrera CP, editor. PLoS One [Internet]. 2021 Aug 12;16(8):e0255226. Available from: https://dx.plos.org/10.1371/journal.pone.0255226

106. Sunyoto T. Partnerships for better neglected disease drug discovery and development: how have we fared? Expert Opin Drug Discov [Internet]. 2020 May 3;15(5):531–7. Available from: https://www.tandfonline.com/doi/full/10.1080/17460441.2020.1736550

